# Massive-scale single-nucleus multi-omics identifies novel rare noncoding drivers of Parkinson’s disease

**DOI:** 10.64898/2026.03.05.709922

**Authors:** Shreya Menon, Adam W. Turner, Serena H. Chang, Alia W. Johnson, Heather H. Chang, Aayushi J. Shah, Youjie Zeng, Colleen E. Strohlein, Lucas Kampman, Courtney Colston, Alexey Kozlenkov, Stella Dracheva, Micol Avenali, Giovanni Palermo, Roberto Ceravolo, Enza Maria Valente, Carolin Gabbert, Joanne Trinh, Geidy E. Serrano, Thomas G. Beach, Global Parkinson’s Genetic Program (GP2), Joshua M. Shulman, Cornelis Blauwendraat, Thomas J. Montine, Zih-Hua Fang, Michael E. Belloy, M. Ryan Corces

## Abstract

Most genetic variants contributing to complex diseases reside in the noncoding genome. While common variants uncovered by genome-wide association studies often fail to explain much of the observed heritability of these diseases, rare variants often have higher effect sizes and cumulatively explain a larger portion of heritability. However, rare variants, particularly rare noncoding variants, have remained under-characterized largely due to the difficulties of accurately predicting variant functionality at scale, given that each individual carries an average of ∼10,000 rare variants. Here, we generated multi-omic data from >3.3 million nuclei sampled from five brain regions across a cohort of 80 individuals with Parkinson’s disease (PD) and 21 neurologically normal control individuals with matched 30x whole-genome sequencing. We use this data to identify cell type-specific features of PD, map cell type-specific chromatin accessibility and expression quantitative trait loci, and train machine learning models to predict the effect of variants on gene regulation. We identify rare noncoding variants statistically associated with sporadic PD and extend our approaches to predict drivers of familial PD of unknown genetic origin. Our results underscore the significance of rare noncoding variants in complex diseases and provide a roadmap for applying similar approaches in other disease systems.

## Introduction

Parkinson’s disease (PD) is an age-related neurodegenerative disease characterized pathologically by the loss of dopaminergic neurons within the substantia nigra as well as the accumulation of Lewy bodies containing abnormal forms of alpha-synuclein in multiple brain regions, and clinically by a constellation of motor and non-motor symptoms^1^. Like many complex diseases, PD is driven by a mix of genetic and environmental factors and is estimated to be ∼30-40% heritable^2,3^. Decades of research have aimed to uncover both the molecular and genetic drivers of PD susceptibility to improve risk stratification and nominate novel targets for therapeutic intervention.

The molecular pathophysiology of PD involves disruption across multiple cell types and brain regions. Recently, single-nucleus transcriptomic profiling (snRNA-seq) has been used to study the cell types and molecular processes most affected in PD. For example, previous work has identified populations of vulnerable dopaminergic neurons marked by *AGTR1* expression^4^ associated with PD-related degeneration and revealed significant glial activation in the midbrain of patients with PD, including microgliosis^5^ and upregulation of stress-induced S100B in oligodendrocytes^5–7^ in the substantia nigra. The pathology of PD extends beyond the midbrain and is associated with widespread microglial activation and epigenetic dysregulation affecting extranigral regions, including the striatum and cortex^8,9^. The involvement of diverse neuronal and glial cell types in PD-related dysfunction underscores the importance of a multi-regional and cell type-specific approach to dissecting PD pathophysiology.

The genetics of PD and nearly all other complex diseases have been studied with two complementary approaches. First, genome-wide association studies (GWASs) have been used to identify common variants, typically present in more than 1% of the population, that are statistically associated with the risk of sporadic P^10–12^. These common variants often have very small effect sizes, with odds ratios frequently between 0.9 and 1.1, and almost universally reside within the noncoding regions of the genome^10^. Noncoding variants can exert a multitude of effects but are thought to primarily act by perturbing gene regulation through alterations to the recognition sites of sequence-specific transcription factors (TFs)^13–15^. However, as these common variants have low effect sizes, they account for less than 10% of the overall risk for PD, leaving much of the heritability of PD unexplained^2,16,17^.

A second approach, familial pedigree analysis, has been used to identify ultra-rare, high-effect-size variants that drive disease. This approach has identified a small number of highly penetrant driver variants in well-studied genes, such as alpha-synuclein (*SNCA*) and parkin (*PRKN*)^2,18^. While highly effective at identifying driver variants within coding regions, these familial pedigrees are often too small to employ classical segregation analyses within the noncoding genome. As 70–80% of familial PD cases lack a clear coding driver variant, this leaves many high-effect-size noncoding variants to be discovered^19–21^. Together, the limitations of existing genetics discovery approaches highlight a need for more robust prioritization of rare noncoding variants.

Rare noncoding variants remain understudied due to challenges related to scale and interpretability. With respect to scale, over 60 million single-nucleotide variants with a frequency less than 0.5% were identified from the 2,504 individuals in the 1000 Genomes Project^22^. These data indicate that, on average, each person has more than 10,000 rare variants, ∼98% of which are noncoding. In fact, ultra-rare variants are estimated to contribute up to 25% of the *cis*-heritability of transcriptional regulation^23^. The abundance of variants and the relatively low number of patients with PD who have whole-genome sequencing (WGS) data available make it challenging to associate rare noncoding variants with disease using existing statistical approaches. Regarding interpretability, the effects of noncoding variants are more challenging to predict than those of coding variants. Gene regulation is highly cell type-specific, and most noncoding variants are not known to exert an impact. We and others^24,25^ have turned to machine learning (ML) approaches to address these problems of both scale and interpretability. These ML models^26^ learn to predict a molecular measurement, such as chromatin accessibility as measured by the assay for transposase-accessible chromatin (ATAC-seq)^27^, from a given sequence. When paired with cell type-specific chromatin accessibility measurements, this ML-based approach enables rapid and robust prediction of the gene regulatory effects of noncoding variants.

Here, we sought to uncover cell type-specific molecular mechanisms of PD and nominate novel genetic drivers of both sporadic and familial disease. We generated a massive-scale single-nucleus multi-omic atlas of the human brain spanning five different brain regions across 80 PD patients and 21 healthy controls. With more than 3.3 million nuclei featuring high-quality, matched snRNA-seq and snATAC-seq data, as well as 30x WGS data for each individual, this dataset provides an ideal resource for understanding PD-specific molecular alterations and training robust ML models for predicting the effects of noncoding variants. We leverage this data to nominate new drivers of the genetic risk of PD, establishing a novel framework for understanding the molecular and genetic underpinnings of complex disease.

## Results

### Single-nucleus multi-omic profiling of multiple brain regions identifies cell type-specific signatures of gene expression and gene regulation

To build a molecular atlas that resolves previously underappreciated rare cell types and aid in the interpretation of noncoding variant effects in PD, we performed paired single-nucleus multi-omic profiling of chromatin accessibility and gene expression from postmortem brain tissue derived from five brain regions across 101 donors (80 PD patients and 21 controls) (**Fig. 1a, Supplementary Fig. 1a**). These individuals were volunteers in the Arizona Study of Aging and Neurodegenerative Disorders community aging study and have detailed clinical and neuropathologic annotation (**Supplementary Fig. 1b–c, Supplementary Table 1**). We selected cases across the neuropathological spectrum of Lewy body formation, which follows a stereotypical progression from brainstem, then limbic, and finally neocortical neurons^28^ (**Supplementary Fig. 1b, Supplementary Table 1**). To mitigate batch effects across this large cohort, we mixed nuclei from different donors together prior to droplet generation and leveraged 30x WGS from each donor to perform genetic demultiplexing (**Supplementary Fig. 1a**). After strict quality control, we created three different atlases: (i) 4.2 million nuclei with high-quality snATAC-seq data, (ii) 4.5 million nuclei with high-quality snRNA-seq data, and (iii) the intersection, 3.3 million nuclei with high-quality multi-omic data (snATAC-seq and snRNA-seq) (**Fig. 1a, Supplementary Fig. 1d–g**). These atlases enabled maximal retention of nuclei for downstream analyses of each modality. In our multi-omic atlas, we retained an average of 32,956 nuclei per individual (across all brain regions), and 92% of the 505 brain samples were represented by over 3,500 nuclei that passed quality control filters in both snATAC-seq and snRNA-seq assays (**Fig. 1b, Supplementary Fig. 2a**).

**Figure 1.**
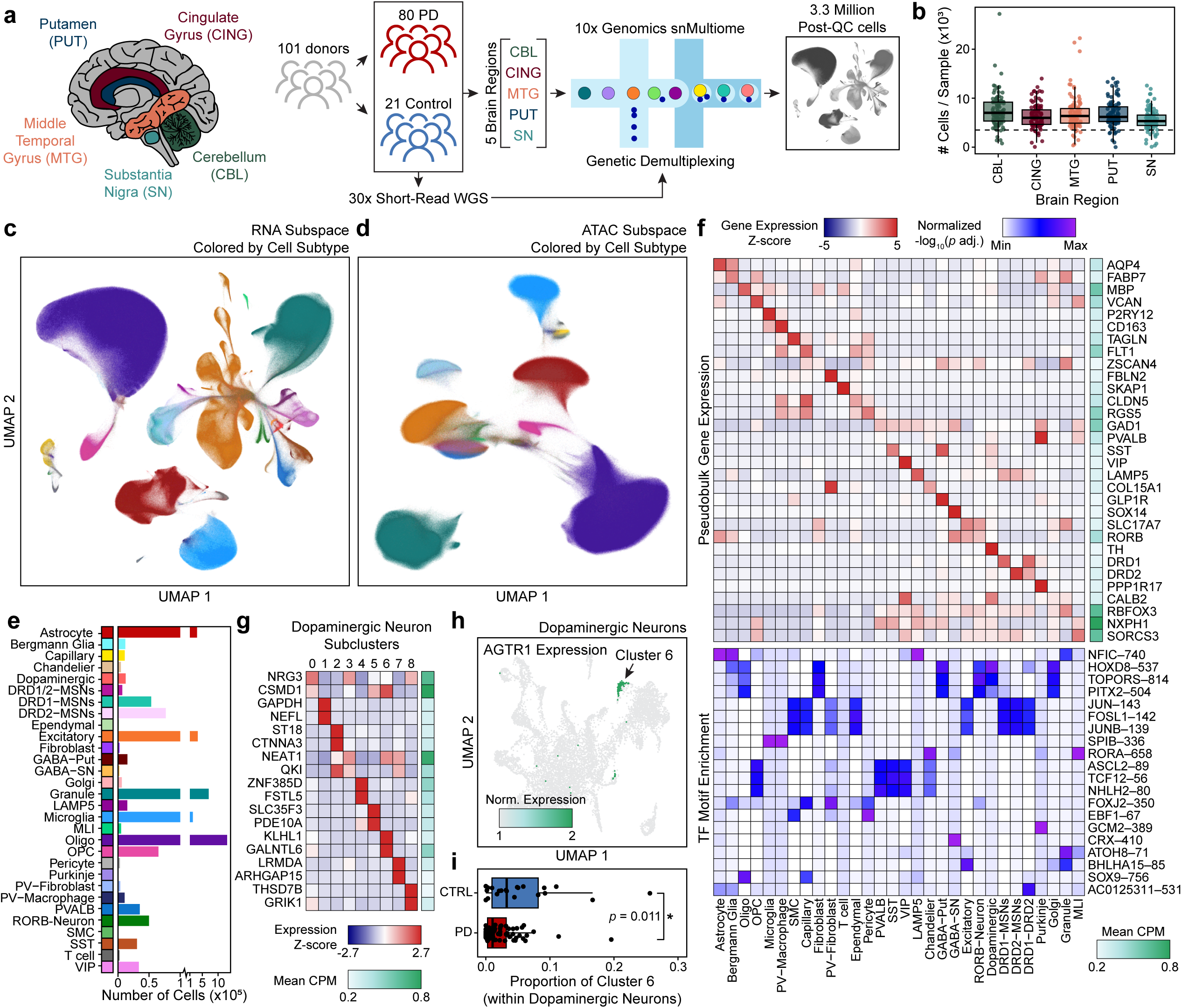
Multi-region single-nucleus multi-omic profiling of postmortem human brain identifies molecular and cellular hallmarks of PD. **a.** Schematic of study design. **b.** Dot plot of the number of post-quality control nuclei recovered per sample, grouped by brain region. Each point represents one sample. Dashed line represents our target recovery of 3,500 nuclei per sample. **c–d.** UMAP embedding of all nuclei based on (**c**) the snRNA-seq modality or (**d**) the snATAC-seq modality, colored by annotated cell subtype. Colors shown in (**e**). **e.** Bar plot of cell counts for each annotated cell subtype across the full dataset. **f.** Heatmap of normalized pseudobulk (top) gene expression and (bottom) transcription factor (TF) motif enrichment across cell subtypes. Rows correspond to genes or motifs; columns correspond to cell subtypes. The right-most column of the top heatmap indicates the average expression in counts per million (CPM) of each gene across all donors in the dataset. **g.** Heatmap showing marker gene expression across dopaminergic neuron subclusters. Columns represent subclusters; rows represent selected genes. The right-most column indicates the average expression in CPM of each gene across all donors in the dataset. **h.** UMAP embedding of all dopaminergic neurons, colored by the scaled log_2_-normalized gene expression values of *AGTR1*. **i.** Box and whisker plot showing the proportion of the *AGTR1*+ dopaminergic neuron subcluster amongst all dopaminergic neurons within each donor, grouped by PD and control samples. The center line denotes the median, the box spans the interquartile range (IQR; 25th–75th percentiles), and the whiskers extend to the most extreme data points within 1.5× the IQR. Each point represents one donor. *p*-value represents results from a two-sided Wilcoxon rank-sum test comparing distributions between groups.

Dimensionality reduction and clustering revealed seven broad cell types (astrocytes, excitatory neurons, inhibitory neurons, microglia/immune cells, oligodendrocytes, oligodendrocyte precursor cells (OPCs), and vascular/endothelial cells), with many neurons showing brain region specificity (**Supplementary Fig. 2b–c**). These cell types showed minimal detectable bias due to covariates, including biological sex, disease status, donor, and batch (**Supplementary Fig. 2d–g**). We further subclustered these broad cell classes based on refined marker genes to identify 30 distinct cell types (**Fig. 1c–f, Supplementary Fig. 3a–d**). To identify which TFs influence cell type-specific regulatory programs, we performed motif enrichment analysis of peaks identified in each cell type (**Fig. 1f**). This analysis identified known cell type-specific master regulators, such as SOX9 in oligodendrocytes^29^ and SPI family TFs in microglia^30^, as well as novel putative regulators, such as GCM2 in Purkinje cells. Notably, motif enrichment analysis also revealed significant enrichment of CRX binding motifs within accessible chromatin regions of GABAergic neurons in the substantia nigra. While CRX has previously been characterized for its role in retinal photoreceptor differentiation^31^, this suggests potential engagement of transcriptional programs not traditionally associated with this neuronal population.

Focusing on subclusters of neurons, we sought to characterize neuronal subtypes in the substantia nigra and putamen, sites of nigro-striatal neuron soma and their afferent synapses with medium spiny neurons (MSNs) that subserve motor functions heavily impacted in PD. Subclustering of dopaminergic neurons revealed transcriptionally distinct populations (**Fig. 1g, Supplementary Fig. 4a–b**). In line with prior work^4^, we observed an *AGTR1*-expressing subcluster (Dopaminergic Subcluster 6) that was reduced in PD compared with controls, providing additional evidence for the selective vulnerability of this dopaminergic neuron population **(Fig. 1h–i**). Additionally, we uncovered substantial evidence for the existence of three distinct populations of MSNs within the putamen (**Supplementary Fig. 4c**). As expected, we identified the canonical MSN subtypes marked by expression of dopamine receptor D1 (*DRD1*) or D2 (*DRD2*) but we also identified a third population that expresses both *DRD1* and *DRD2* in addition to other previously unappreciated marker genes (**Supplementary Fig. 4d–f**). This population corresponds to “eccentric” MSNs^32,33^, a transcriptionally distinct subtype of MSNs that lies outside the canonical dichotomy of striatal output pathways. Intriguingly, DRD1-and DRD2-MSNs separate more cleanly in the RNA subspace than in the ATAC subspace, while the DRD1/2 MSNs remain clearly delineated in both.

We further subclustered the most abundant non-neuronal cell types, uncovering varying degrees of regional specificity (**Supplementary Fig. 5a–b**). Differential expression analysis identified subpopulation-specific markers, such as *KLHDC7A*+ astrocytes in the substantia nigra (Astrocyte Subcluster 1) and *IGLC1*+ microglia in the substantia nigra (Microglia Subcluster 2), suggesting that non-neuronal cells adopt regionally specialized transcriptional programs tailored to local environments (**Supplementary Fig. 5c**).

### Cell type-specific transcriptional and cellular alterations reveal coordinated glial and neuronal responses in PD

Our dataset provides three primary angles from which to understand the molecular pathogenesis of PD: (i) comparison of gene expression and chromatin accessibility differences between cases and controls, (ii) comparison of cellular proportions between cases and controls, and (iii) comparison of gene expression across the distribution of Lewy bodies within the brain.

Using a case-control paradigm, we first performed differential gene expression analysis between cases (N = 80) and controls (N = 21) using pseudobulked expression profiles from each individual within each cell type (**Supplementary Table 2**). For each test, we corrected for age at death, biological sex, and postmortem interval if those covariates showed a correlation (Pearson’s *r p*-value < 0.05) with any of the first five principal component values for pseudobulked expression of that cell type (**Supplementary Fig. 6a**). We also performed a permutation-based assessment of the reliability of those differentially expressed genes (DEGs) by shuffling the case/control labels and determining the background number of DEGs that would occur by chance within our dataset, an approach that explicitly accounts for the imbalanced sample sizes of PD and control individuals in our cohort (**Supplementary Fig. 6b**). This analysis highlighted oligodendrocytes, MSNs, excitatory neurons, and astrocytes as the cell types with the greatest changes in transcriptional profile (**Fig. 2a, Supplementary Fig. 6c**). We note that, as expected, the total number of DEGs detected is impacted by the number of cells of the given cell type captured per sample and thus some cell types may remain underpowered even in this large dataset (**Supplementary Fig. 6D**). Although dopaminergic neurons did not pass permutation-based thresholds, pathway analysis of the genes identified converged on dopamine uptake and catecholamine transport. Excitatory neurons similarly showed enrichment for monoamine and catecholamine response pathways, together implicating dysregulated neurotransmitter signaling as a central mechanism in PD pathogenesis^34^ (**Supplementary Fig. 7a, Supplementary Table 3**). To further understand regulatory changes associated with transcriptional dysregulation, we performed a targeted analysis of chromatin accessibility at regulatory elements mapping to DEGs, identifying gene-associated peaks with differential accessibility between cases and controls (**Supplementary Table 4**). Regulatory concordance between chromatin accessibility and gene expression was most pronounced in MSNs, where over half of the DEGs were associated with at least one differentially accessible regulatory element, whereas glial cell types exhibited fewer detectable accessibility changes at DEG-linked regions.

**Figure 2.**
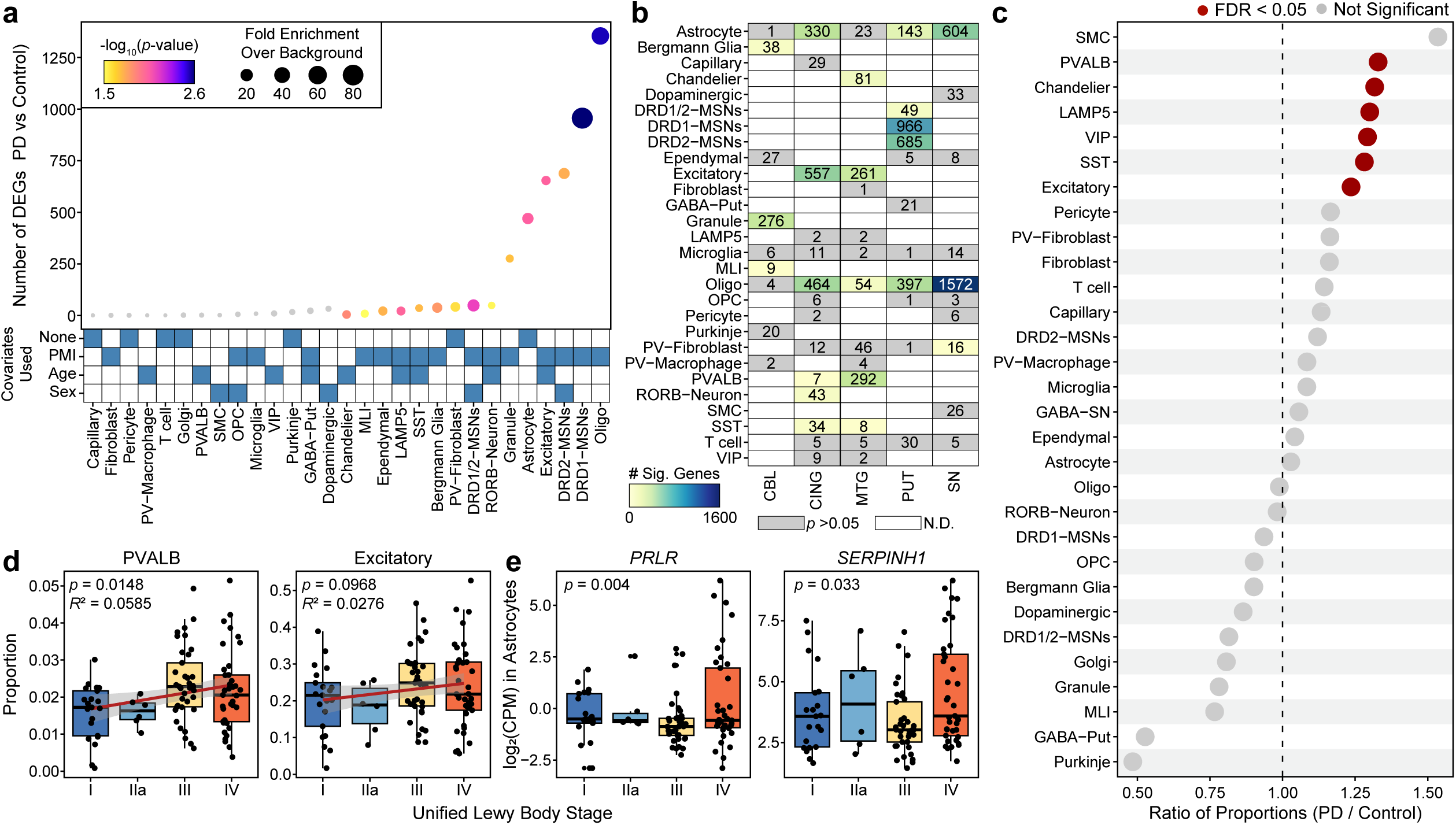
Distinct cell types show differential gene expression and proportion differences in PD. **a.** Dot plot showing the number of differentially expressed genes (DEGs) identified in each cell type. Point size represents the fold enrichment of the number of DEGs relative to a permuted background, and point color represents the –log_10_ of the empirical *p*-value of the number of DEGs calculated from permutation analysis. Points shown in gray color did not pass the permutation test significance threshold. The heatmap below indicates which covariates (none, postmortem interval (PMI), age at death, and/or biological sex) were included for each cell type based on correlation with principal components. **b.** Heatmap showing the number of significant DEGs per cell type, separated by brain region. Cells are shaded by the count of significant DEGs; gray indicates a permutation-based *p-*value > 0.05; white indicates not determined due to the non-existence of that cellular subtype in the specified brain region. **c.** Dot plot showing the ratio of cell type proportions in PD relative to control. Each point represents one cell type; red color indicates FDR < 0.05, gray indicates not significant. The dashed line marks a ratio of 1. **d–e.** Box and whisker plots showing (**d**) the proportion of (left) PVALB and (right) Excitatory neuron populations or (**e**) log_2_(counts per million (CPM) pseudobulk expression) of (left) *PRLR* and (right) *SERPINH1* of astrocytes across stages of the of the Unified Staging System for Lewy Body Disorders. Per box and whisker plot, the center line denotes the median, the box spans the interquartile range (IQR; 25th–75th percentiles), and the whiskers extend to the most extreme data points within 1.5× the IQR. Each point represents a donor. In (**d**), the *p*-values reflect likelihood ratio tests from linear regression models, evaluating whether cell type proportions vary with Lewy body stage. In (**e**), the *p*-values reflect likelihood ratio tests evaluating whether gene expression varies with Lewy body stage, adjusting for disease status.

When further stratifying this analysis based on brain region, we found that the most drastic changes in oligodendrocytes and astrocytes occurred within the substantia nigra, the site of the most pronounced neurodegeneration in PD (**Fig. 2b**). Across astrocytes, oligodendrocytes, and excitatory neurons, these DEGs showed greater overlap than expected by chance between brain regions based on permutation testing (**Fig. 2b, Supplementary Fig. 7b**), suggesting that a subset of transcriptional responses in PD is coordinated across regions and reflecting shared injury or response to injury mechanisms. Specifically, among oligodendrocytes, the strongest overlap occurred between the substantia nigra and putamen (**Supplementary Fig. 7c–d**). Enrichment analysis further revealed that Oligodendrocyte Subclusters 0 and 1 were most strongly enriched for regional DEGs (**Supplementary Fig. 7e**). The broad regional representation in Oligodendrocyte Subcluster 0 suggests a common oligodendrocyte response to PD, whereas the overrepresentation of substantia nigra oligodendrocytes in Oligodendrocyte Subcluster 1 points to region-specific injury or response to injury in this prominent site of neurodegeneration. Pathway analysis of these overlapping regional DEG sets highlights nitric oxide–cGMP signaling, perisynaptic extracellular matrix organization, and non-integrin membrane-extracellular matrix interactions, pointing to shared alterations in synaptic plasticity and neuron–glia communication across these disease-relevant brain regions **(Supplementary Fig. 7f)**. Together, these findings implicate extracellular and intracellular mechanisms in shaping oligodendrocyte gene expression changes within the nigrostriatum. As expected, the cerebellum, a region largely spared from neurodegenerative features of PD, showed comparatively fewer significant gene expression changes (**Fig. 2b**). Overall, these molecular alterations—particularly those observed in oligodendrocytes—are consistent with previous reports in PD^35–38^ and underscore the coordinated molecular responses across cell types and regions within the diseased brain.

In contrast to these differential expression results, we identified a largely distinct set of cell types that showed significant differences in cellular proportions between PD cases and controls. Whereas the majority of our differential expression results were identified in glial cells, all of the cell populations that showed significant differences in cellular proportions were neuronal, including multiple types of inhibitory neurons showing increased proportion in PD (**Fig. 2c, Supplementary Fig. 8a**). This could be explained by compensatory mechanisms in glial cells in response to neurodegeneration or could reflect the loss of some neuron types in response to molecular alterations occurring in glial cells. Within the two brain regions profiled that contain these neuronal types, the cellular proportion differences were more pronounced in the cingulate gyrus than in the middle temporal gyrus (**Supplementary Fig. 8b**). The prominence of changes in the cingulate gyrus is consistent with prior neuropathological and imaging studies showing early Lewy body deposition and neuronal loss in this region, features that have been associated with cognitive decline and neuropsychiatric symptoms in individuals with PD^35,39,40^.

To better understand how these changes in transcription and cellular proportions relate to the neuropathological progression of PD, we stratified our cohort based on the Unified Staging System for Lewy Body Disorders^41^ that has Stage I (N = 21) for olfactory bulb or amygdala Lewy bodies, which are not neuropathological features of PD progression, and subsequent stages that reflect the stereotypical progression of Lewy body accumulation in individuals with PD: brainstem (Stage IIa; N = 6), then limbic (Stage III; N = 37), and finally neocortical (Stage IV; N = 36) neurons. Stratifying across this progression, we observe that the same cell types showing changes in cell proportion between disease and control in the cingulate gyrus also show progressive changes across disease stages (**Fig. 2d and Supplementary Fig. 8c**). At the transcriptional level, several genes displayed stage-associated expression dynamics. Notably, *SERPINH1* (aka *HSP47*), a gene encoding a collagen-specific chaperone implicated in proteostasis and neuroinflammation in PD^7,42^, and *PRLR*, a gene encoding the prolactin receptor which is linked to dopaminergic signaling^43^, both demonstrated increased expression in astrocytes at late disease stages (**Fig. 2e**). We also identified chromatin accessibility peaks from astrocytes that mapped to *SERPINH1* that showed stage-associated changes, suggesting that its transcriptional upregulation may be driven by dynamic regulatory mechanisms (**Supplementary Fig. 8d**). To further resolve cell type-specific stage-associated transcriptional patterns, we classified these genes based on their trajectories through the pathological progression of PD (**Supplementary Fig. 8e, Supplementary Table 5–6**). MSNs were particularly enriched for genes showing monotonic upregulation across Lewy body stages, highlighting progressive, stage-linked transcriptional activation in this neuronal population (**Supplementary Fig. 8f**). Overall, these findings reveal progressive alterations in both cellular state and cellular composition, pointing to mechanisms that may underlie the vulnerability of specific cell types in PD.

### Paired WGS and molecular measurements identify cell type-specific expression and chromatin accessibility quantitative trait loci

Our dataset of snRNA-seq and snATAC-seq paired with WGS is an ideal resource from which to nominate cell type-specific quantitative trait loci (QTLs) that would inform common variant associations with brain-related traits and diseases. We called chromatin accessibility QTLs (caQTLs) using RASQUAL^44^ on the 4.2 million nuclei snATAC-seq atlas and called expression QTLs (eQTLs) using TensorQTL^45^ on the 4.5 million nuclei snRNA-seq atlas (**Supplementary Data 1**). In each cell type, we limited our analyses to individuals where at least 50 nuclei were available. As expected, the number of QTLs called generally increased with the number of cells available for each cell type (**Fig. 3a**). On average, 50% of the cell type-specific eQTLs we identified were novel compared to the tissue-level eQTLs called in the Genotype-Tissue Expression (GTEx) project^46^ (**Fig. 3b–c**). Many of the eQTLs that we identified were unique to a single cell type, and these were particularly poorly represented in GTEx, especially for less abundant neuronal subtypes such as dopaminergic neurons (**Fig. 3d**). Similarly, our caQTL analysis revealed numerous cell type-specific regulatory variants, several of which overlapped loci implicated by GWAS. As an illustrative example, we identified a SNP (rs72836333) that maps to a regulatory peak predicted to interact with *KANSL1* (**Fig. 3e**), a gene located within the *MAPT* locus and implicated in neurodegenerative disease risk^47,48^. rs72836333 is in strong linkage disequilibrium (r^2^ > 0.99) with the PD-associated lead variant rs113564729^49^, indicating that it tags the same disease-associated haplotype. Further, rs72836333 is an eQTL in GTEx for multiple brain regions, and the A allele is predicted to create a TGIF1 TF motif (**Fig. 3e**).

**Figure 3.**
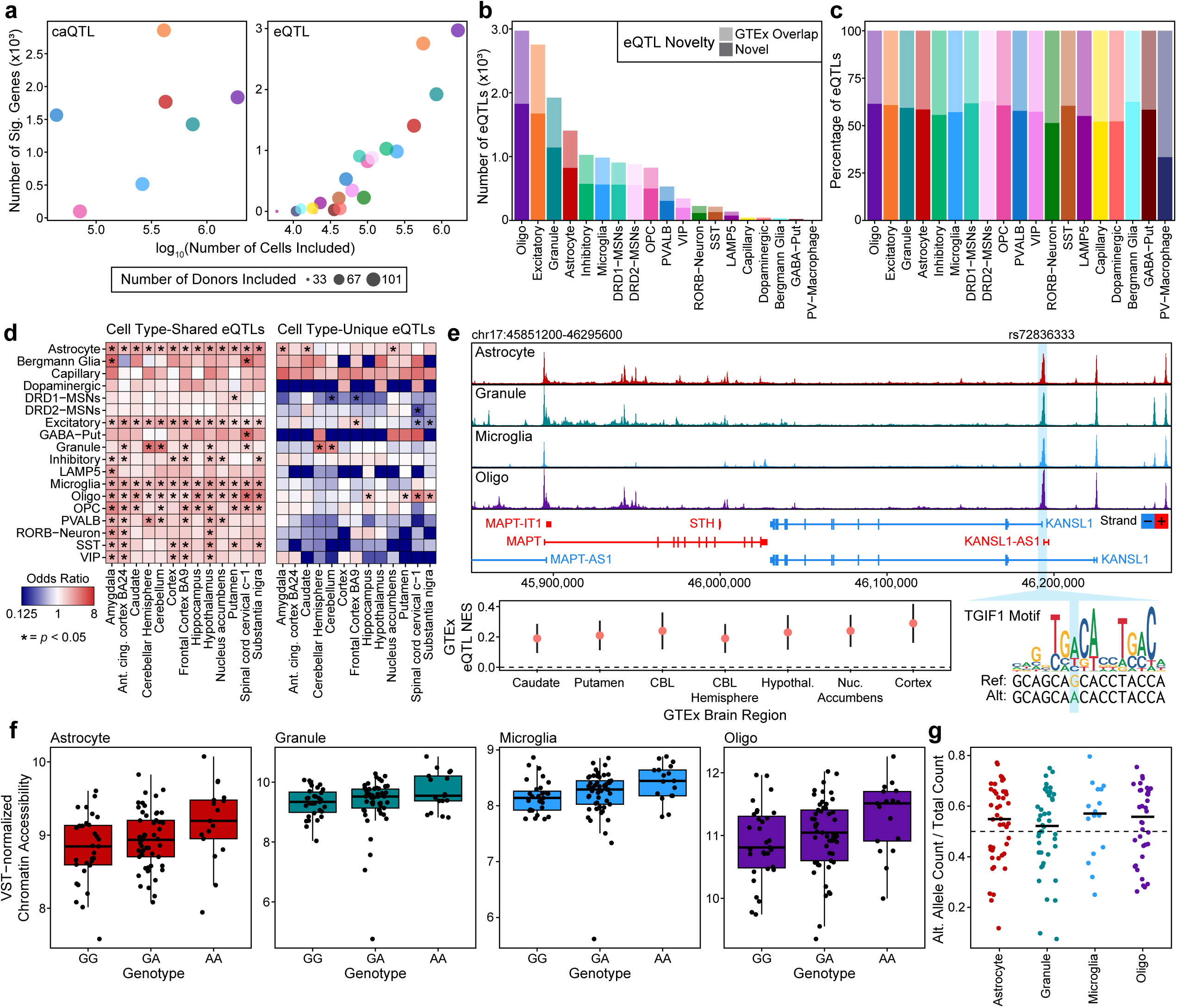
Cell type-resolved cis-regulatory QTL mapping across cell populations identify functional common noncoding variants. **a.** Dot plot showing the number of significant (left) caQTLs and (right) eQTLs detected per cell type as a function of the number of cells included. Each point and color represents a cell type (as shown in **b**). Point size represents number of donors included in QTL testing. **b–c.** Bar plot showing (**b**) the number and (**c**) the percentage of eQTLs detected in each cell type, separated into those overlapping GTEx eQTLs (lighter) and those not detected in GTEx (darker). **d.** Heatmap of odds ratios for the overlap of QTLs identified in each cell type (y-axis) and GTEx region (x-axis), split between cell type-shared QTLs indicating those found in >1 cell type (left) and cell type unique (right). Odds ratios and *p*-values were computed using a two-sided Fisher’s exact test, with *p*-values adjusted for multiple testing using the Benjamini-Hochberg method. Asterisks indicate FDR-adjusted *p* < 0.05. **e.** ATAC-seq tracks showing normalized pseudobulked chromatin accessibility signal across selected cell types in the rs72836333 QTL locus near *MAPT/KANSL1*. Bottom left, effect sizes represented as the log_2_(allelic fold change) are shown for GTEx brain regions at the corresponding eQTL; points represent different GTEx tissues. Error bars indicate 95% confidence interval of effect size. Bottom right, predicted affected TF motif of TGIF1. NES = Normalized Enrichment Score. **f.** Box and whisker plots showing variance-stabilized chromatin accessibility values stratified by genotype for rs72836333 in the indicated cell types. The center line denotes the median, the box spans the interquartile range (IQR; 25th–75th percentiles), and the whiskers extend to the most extreme data points within 1.5× the IQR. Each point represents an individual donor. **g.** Scatter plot showing reference and alternate allele read counts for rs72836333 across pseudobulk donors, colored by cell type. Each point represents one donor–cell type pseudobulk. The dashed line denotes equal reference and alternate counts.

Within our atlas, rs72836333 functions as a caQTL in astrocytes, granule cells, microglia, and oligodendrocytes (**Fig. 3f**), and we find evidence of allelic imbalance in each of those cell types within our snATAC-seq data (**Fig. 3g**).

### Machine learning models enable rapid prediction of noncoding variant effects

To enable high-throughput prioritization of noncoding variant effects, we trained convolutional neural network models to predict chromatin accessibility from DNA sequence. Specifically, we used ChromBPNet^50^ to train cell type-specific models from the 25 cell types with sufficient representation in our dataset. We trained these ML models exclusively on the 21 control individuals to avoid data leakage from genetic signals from patients with PD (**Fig. 4a**). We then used these models to prioritize all rare noncoding variants observed in the 3,611 patients with PD represented in the Global Parkinson’s Genetics Program (GP2; **Fig. 4a**), which includes the 80 patients with PD sequenced in our study. Here, we define rare noncoding variants as those variants that reside within a peak region in our dataset and have an allele frequency below 0.01 in all populations in the Genome Aggregation Database (gnomAD)^51^. We quantified the ML-predicted impact of each variant using the Jensen–Shannon Divergence (JSD) between the predicted chromatin accessibility profile of the reference and alternate alleles, allowing us to create a prioritized list of variants predicted to increase or decrease chromatin accessibility (**Fig. 4b–c, Supplementary Fig. 9a**). These 25 ML models each showed robust performance in predicting chromatin accessibility on held-out data (avg. Pearson’s *r* = 0.73; range 0.68–0.82), with no clear relationship between model performance and number of cells used for training (**Supplementary Fig. 9b**). This high performance was observed both in genome-wide peak regions and in cell type-specific peak regions, underscoring a key advantage of using cell type-specific models over composite models of chromatin accessibility (**Fig. 4d**). In each cell type, we tested only the variants found within peak regions of the cell type, and on average, prioritized 4% of all tested variants per cell type (**Fig. 4e**). The vast majority of variants had predicted effects only in a single cell type (**Fig. 4f**), consistent with the well-known cell type-specificity of gene regulation^52^. Similarly, we observed a significant enrichment of rare variants among the top 1% of effect scores compared with common variants (**Supplementary Data 2**, **Fig. 4g**, Fisher’s exact test, *p*-value < 2.2×10^−16^; OR = 1.23; 95% CI: 1.22–1.24). Additionally, rare variants exhibited significantly higher effect scores overall compared to common variants (Wilcoxon rank-sum test, *p*-value < 2.2×10^−16^), supporting the observation that rarer alleles are more likely to exert strong predicted effects. As expected, given the distribution of peak regions, we further found that most prioritized variants reside in distal/intergenic or intronic regions (**Fig. 4h**). In all, we performed a total of 88,611,830 predictions across 25 cell types and prioritized 3,245,735 variants for further analysis (**Supplementary Data 2**).

**Figure 4.**
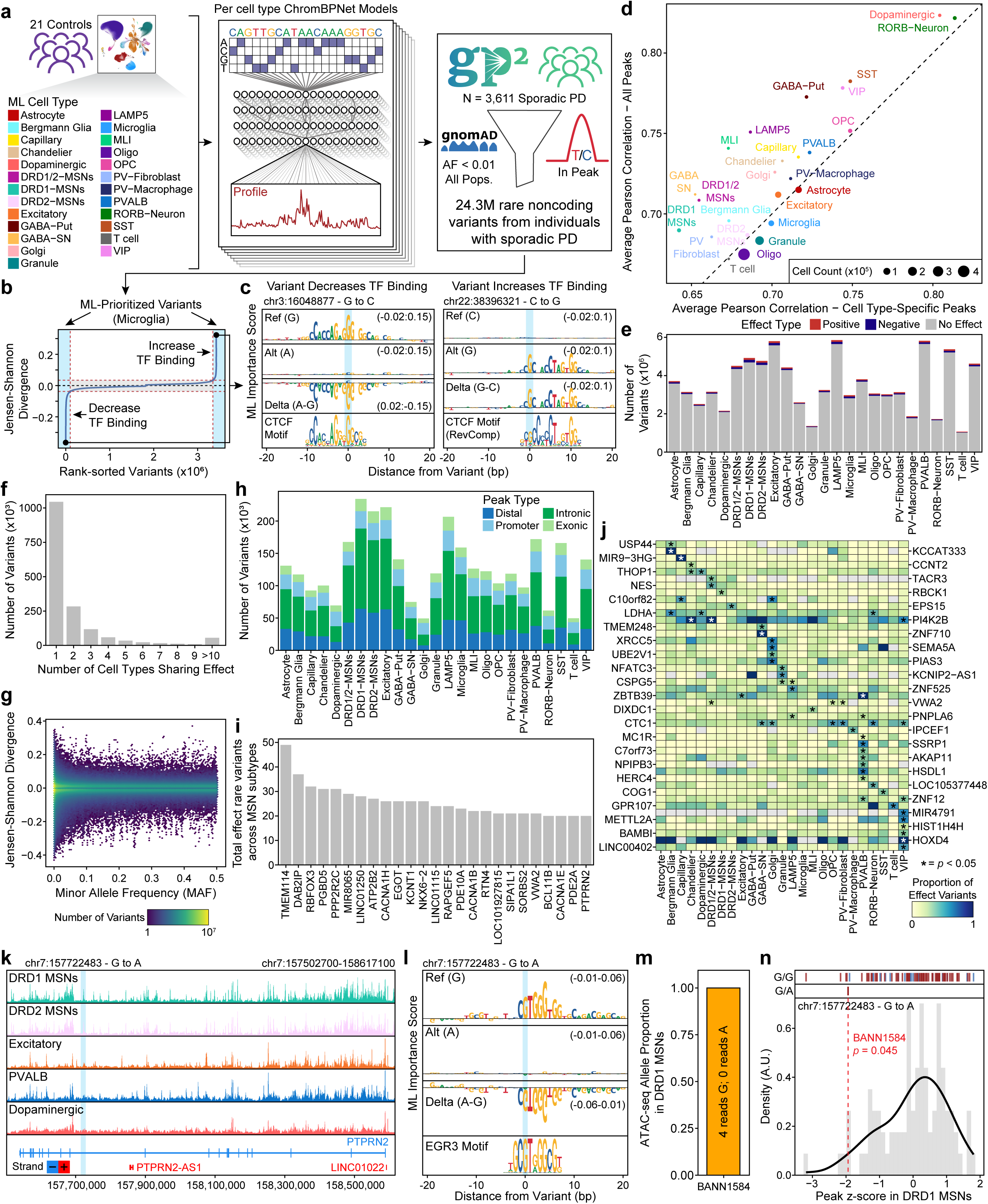
Modeling cell type-specific chromatin effects highlights putatively functional rare noncoding variants found in sporadic PD. **a.** Schematic of the noncoding variant effect prediction workflow for sporadic PD. **b.** Representative distribution plot of ML-based scores for all rare noncoding variants in microglia. The inflection points (red dashed lines) of the genome-wide score distribution (Jensen–Shannon Divergence) defines variants predicted to increase or decrease TF binding. **c.** Importance score plots of predicted TF binding changes at (left) the bottom scoring and (right) top scoring variants. The relative size of the DNA base within the logo represents the position’s importance. Importance score ranges for each plot are shown in parentheses. **d.** Scatterplot comparing ChromBPNet performance across cell types. Model performance is defined as the average Pearson’s *r* between observed and predicted pseudobulk ATAC-seq counts, averaged across 5 cross-validation folds. The y-axis shows performance across all peaks, and the x-axis shows performance across peaks specific to each cell type. Point size reflects the number of cells used to train the model. **e.** Stacked bar plot showing counts of variants predicted to have positive (red), negative (blue), or no effect (grey) per cell type. **f.** Bar plot showing the number of variants as a function of the number of cell types in which each variant is predicted to have an effect. **g.** Hexbin plot colored by point density of variant effect scores versus minor allele frequency for all variants across all cell types. **h.** Bar plot showing the count of variant genomic annotations across cell types, colored by the peak type. **i.** Bar plot showing the number of distinct effect variants observed per gene in each MSN subtype (DRD1-, DRD2-, and DRD1/2-MSNs), summed across subtypes (variant–subtype pairs). Genes are ranked by this summed effect burden, and the top 25 are shown. **j.** Heatmap showing the proportion of all variants attributed to each gene that are predicted to have an effect by ML. Proportions are shown across cell types for genes identified as enriched for effect variants. Rows represent genes and columns represent cell types. Asterisks denote genes significantly enriched for effect variants based on permutation testing. **k.** Normalized pseudobulked chromatin accessibility tracks for the genomic region surrounding *PTPRN2*. The area immediately surrounding variant chr7:157722483 - G to A is highlighted in light blue. **l.** Importance score plots for the variant shown in (**k**), including reference, alternate, and delta signal, with the associated EGR3 TF motif shown below. Importance score ranges for each plot are shown in parentheses. **m.** Bar plot showing the proportion of ATAC-seq reads with the corresponding allele (yellow – G) from DRD1 MSNs cells identified from BANN1584, the individual harboring the variant shown in (**k**). **n.** Density plot of z-score-normalized accessibility for the variant-overlapping peak in DRD1 MSNs for individuals with non-zero accessibility. The dashed red line indicates the accessibility of BANN1584 who harbors the corresponding variant shown in (**k**). Genotype is shown above as tick marks, and the tick mark color denotes disease status (red – PD, blue – Control).

### Mapping of putative enhancer-gene interactions using chromatin conformation capture and single-nucleus multi-omics

With these prioritized variants in hand, we next sought to map putative regulatory elements to their target genes with the goal of predicting the functional consequences of noncoding variants in our data. To do so, we generated cell type-specific chromatin conformation data (Micro-C) from neurons, oligodendrocytes, microglia, and astrocytes isolated by fluorescence-activated nuclei sorting of middle temporal gyrus tissue (**Supplementary Fig. 10a–c**). To obtain sufficient sorted nuclei from each cell type, we pooled nuclei across many donors prior to sorting (see **Methods**). Given the high depth of the data, we called “loops”^53^ at 10-kb resolution, identifying 120,512 total loops (**Supplementary Fig. 10d–e**). As expected, the majority of loops spanned distances less than 500 kb, with 57–70% of anchors overlapping with a called ATAC-seq peak in the respective cell type (**Supplementary Fig. 10f–l**).

We then employed a tiered approach to link regulatory elements to their putative target genes (**Supplementary Fig. 11a**). First, we used the Activity-By-Contact (ABC) model^54^, which predicts enhancer-gene connections based on chromatin accessibility and three-dimensional genome architecture. These ABC models were constructed using our cell type-specific snATAC-seq and Micro-C data, along with publicly available cell type-averaged Hi-C data and cell type-specific H3K27ac ChIP-seq when applicable. For peaks without ABC mapping, we used correlation-based peak-to-gene links^55^, selecting associations with strong correlation scores (*r* ≥ 0.4) and statistical significance (FDR ≤ 0.01). When multiple linked peaks overlapped with a query peak, we prioritized the peak with the greatest physical overlap and selected the gene with the strongest correlation to the query peak. Finally, for the remaining unmapped peaks, we assigned the nearest gene within 25 kb of the transcription start site as a potential target. Any remaining unmapped peaks and the variants they contained were excluded from downstream analyses. This hierarchical strategy yielded comprehensive regulatory element annotations while prioritizing evidence-based functional connections over simple proximity-based assignments (**Supplementary Fig. 11b–d**).

### Rare noncoding variants impact PD-relevant genes

Application of this prioritization strategy highlighted biologically coherent sets of genes with disproportionate burdens of predicted regulatory variation. Given the central relevance of MSN subtypes highlighted throughout our work, we next focused on genes with high regulatory burden in MSNs, identifying the genes with the highest summed effect variant burden across MSN subtypes, including known neuronal regulators such as *RBFOX3*, *PDE10A*, and *PTPRN2* (**Fig. 4i**). To extend this analysis across all cell types, we ranked genes by their normalized burden of rare noncoding effect variants, accounting for differences in the number of regulatory elements mapped to each gene. This revealed several genes involved in protein trafficking and the ubiquitin–proteasome system, including *KLHL12*, *TVP23B*, and *LRRC59*, as well as *THOP1*, which contributes to peptide turnover (**Supplementary Fig. 11e**). To assess whether specific genes concentrate their effect variants within particular cell types more than expected by chance, we performed a permutation-based test of cell type-specificity. For each gene, we computed two complementary metrics: (i) the top-share, representing the maximum proportion of effect variants in any single cell type, and (ii) the τ (tau) index, which quantifies the overall degree of cell type restriction across all cell types. We shuffled cell type labels among effect variants within each gene to generate empirical null distributions, then identified genes whose observed top-share or τ exceeded the 95th percentile cutoffs. This analysis revealed that 3,230 of 13,944 genes (23.2%; 95% CI, 22.5–23.9%) exhibited significant cell type-restricted enrichment of effect variants (**Supplementary Fig. 11f**), indicating that approximately one-fifth of genes concentrate their regulatory variation within specific cell types which is substantially higher than would be expected if effect variants were randomly distributed across cell types. This is consistent with the hypothesis that a number of rare regulatory variants exert effects through highly localized transcriptional programs.

While this analysis identifies genes with cell type-specific *patterns* of effect variants, it does not directly assess whether individual cell types harbor disproportionate burdens of effect variants in particular genes. To identify which genes showed enrichment of effect variants in specific cell types, we performed a complementary permutation-based analysis. For each gene in each cell type, we aggregated the total number of unique effect variants and compared the observed count to a null distribution generated by shuffling cell type labels within each gene. This approach identified genes with significantly elevated effect variant burdens in particular cell types, with the majority showing enrichment in only a single cell type (**Supplementary Fig. 11g**). Notably, we identified a significant enrichment of effect variants in *LDHA* within dopaminergic neurons (**Fig. 4j**). Given *LDHA*’s central role in lactate metabolism and energy homeostasis, this finding suggests that genetic disruptions to metabolic processes may contribute to the selective vulnerability of dopaminergic neurons in PD. Across all tested cell types, genes enriched for cell type-specific effect variants converged on pathways regulating protein depolymerization, ubiquitination, and DNA repair, highlighting mechanisms of proteostasis and genome integrity that are increasingly implicated in PD pathogenesis (**Supplementary Fig. 11h**).

As an illustrative anecdote, we also identified a rare noncoding variant mapping to *PTPRN2* (chr7:157722483 - G to A; **Fig. 4k**), which regulates vesicle-mediated secretion and has been implicated in epigenetic regulation in metabolic diseases^56,57^. This variant resides within an intronic putative regulatory element of *PTPRN2* in both DRD1 and DRD2 MSNs and is predicted to disrupt an EGR3 TF motif based on our ChromBPNet models of those cell types (**Fig. 4l**). This variant was identified within one of the 80 PD individuals within our cohort, enabling us to use the snMultiome data to observe whether the variant indeed decreases TF binding at this site in MSNs. Though the total depth at this site is expectedly low due to sparsity, from 211 DRD1 MSNs we recovered four reads, all from the G allele (**Fig. 4m**). This allelic imbalance matches the expected direction, supporting the predicted regulatory impact of the alternate allele. Moreover, amongst individuals with non-zero accessibility in our multi-omic cohort, this individual was in the bottom 6th percentile for depth-normalized chromatin accessibility of the corresponding peak region (**Fig. 4n**), further confirming the expected effect. Another representative example identified through this prioritization schema is a rare noncoding variant (chr8:3335995 G to A) within an intronic region of *CSMD1*, a complement-regulating gene previously linked to familial PD^58^. The variant, observed in an individual from the GP2 cohort, is predicted to reduce chromatin accessibility in GABAergic interneurons in the putamen and in excitatory and SST+ interneurons across the brain by disrupting a JUNB-binding motif (**Supplementary Fig. 11i–j**). Collectively, these findings underscore the ability of ML-informed prioritization to uncover both pathway-level convergence and cell type-specific signals from rare noncoding variation in PD.

### ML-prioritization enables rare noncoding variant association testing in PD

While the analyses described above provide insights into which genes are frequently disrupted by rare noncoding variants in PD, they do not statistically associate the disruption of those genes with the development of PD; this requires the use of rare variant association testing. However, rare variant association testing within the noncoding genome is typically hindered by the large number of rare noncoding variants and the limited availability of whole-genome sequences. To identify novel genes statistically associated with PD through rare noncoding variants, we developed an ML-informed framework for rare variant association testing (**Fig. 5a**). Using our snATAC-seq-defined putative regulatory elements and the previously described mapping of regulatory elements to target genes, we first defined “testing units” for each cell type. A single testing unit corresponds to all of the regulatory elements mapped to an individual gene, allowing us to compile signals across regulatory elements and reduce the total number of tests performed. Using ML-prioritized variants within these testing sets from the 3,611 PD cases and 350 controls available in the GP2 Release 8, we performed rare variant association tests using SKAT^59^, weighting each variant by its allele frequency and its ML-predicted effect (JSD) with well-calibrated test statistics across cell types (genomic inflation factor λ = 0.80-0.87, **Supplementary Fig. 12a**). On average, we tested associations for 14,773 genes per cell type (**Supplementary Fig. 12b**), identifying 46 genes with statistically significant associations within at least one cell type (**Fig. 5b**, large dots; **Supplementary Table 7**). After multiple hypothesis correction across all cell type-specific tests, 16 gene associations remained globally significant (**Fig. 5b**, bolded gene names). Some of the cell type-specific hits target pathways of known relevance to PD. For example, we identified PD-associated rare noncoding variants regulating the *FAHD1* gene that are associated with PD in astrocytes, oligodendrocytes, and Bergmann glia. *FAHD1* converts enol-oxaloacetate, a strong inhibitor of succinate dehydrogenase, to oxaloacetate, thus improving aerobic respiration efficiency by preventing succinate dehydrogenase inhibition^60^. Other hits target emerging mechanisms related to PD, such as the association observed with *DICER1* in microglia, which could be relevant to recent reports citing transfer RNA fragments as a putative biomarker in PD^61^. Moreover, there were some associations that did not withstand cross-cell type multiple hypothesis correction but nonetheless pointed to intriguing biological mechanisms. For example, rare noncoding variants in dopaminergic neurons (and other cell types) linked to the *PHYHD1* gene which codes for an enzyme that catalyzes the conversion of 2-oxoglutarate to succinate and whose dysregulation could impair mitochondrial respiration (**Fig. 5b**). However, most hits point to potentially novel cellular mechanisms of disease, necessitating confirmatory studies.

**Figure 5.**
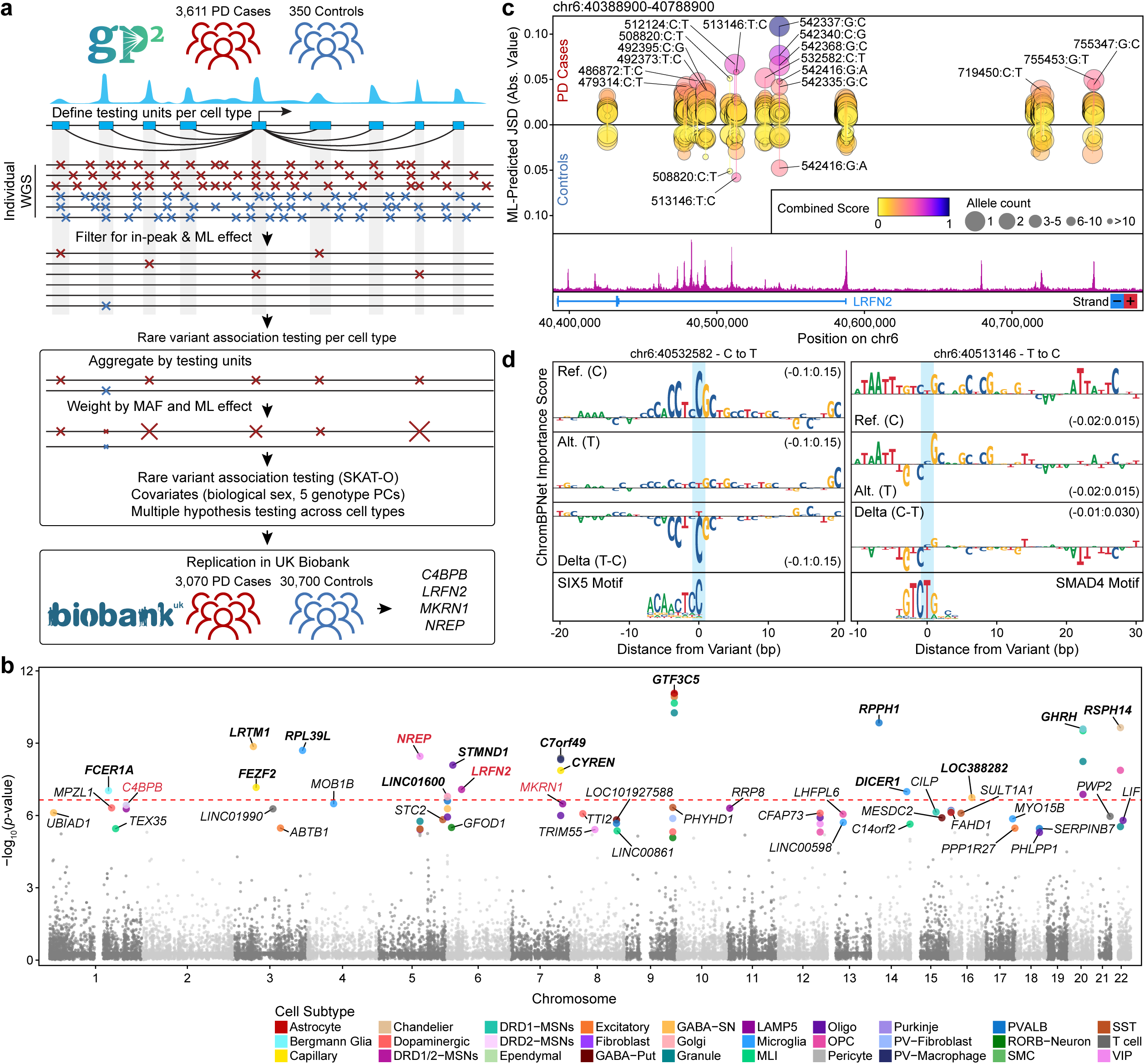
Integrating cell type-specific regulatory predictions with rare variant association testing identifies PD-associated loci. **a.** Schematic of the rare variant association testing workflow. **b.** Manhattan plot showing –log_10_(*p*-value) from cell type-specific rare variant association tests using mapped genes as testing units. Loci passing the significance threshold per cell type are labeled and have larger points colored by the cell type in which the association was detected. Genes passing a global significance threshold accounting for all tests made across cell types are labeled by bold text. Genes that were replicated in the UK BioBank cohort are labeled in red. **c.** Lollipop plot of identified variants in the *LRFN2* locus within the GP2 cohort. Variants are shown separately for PD cases and controls. The x-axis denotes genomic position, and the y-axis shows the absolute value of the predicted effect score (Jensen–Shannon Divergence) from the DRD1/2 ML model. Points are colored by the combined score of minor allele frequency and ML-predicted effect, point size reflects allele count, and the text label represents the last 6 digits of the chromosomal position of the variant in the form of “40,###,###”. The track below shows normalized pseudobulk chromatin accessibility for DRD1/2 MSNs across the same region, aggregated across all donors. **d.** Predicted TF binding changes for representative variants at the *LRFN2* locus from DRD1/2 ML models. Reference and alternate allele importance tracks are shown with the corresponding delta signal. Motif logos for the affected (left) SIX5 and (right) SMAD4 motifs are shown below. Importance score ranges for each plot are shown in parentheses.

To replicate these findings in an unrelated cohort, we curated a set of 3,070 individuals with a clinical diagnosis of PD and 30,700 sex-matched controls from UK Biobank whole-genome sequencing data following stringent ancestry, quality control, and relatedness filters. We selectively searched all cell type-significant positive testing units from our initial cohort for PD association within the UK Biobank. Of the initial 78 cell type-significant associations, four remained significant in the validation cohort (*LRFN2* in DRD1/2 MSNs, *C4BPB* in DRD1 MSNs, *MKRN1* in LAMP5 neurons, and *NREP* in Chandelier cells) (**Fig. 5b**, red gene names; **Supplementary Fig. 12c**). While this replication rate is low, it is in line with the comparatively small size and low number of control individuals in the discovery cohort. The association of variants in *LRFN2* with MSNs is particularly interesting given the known importance of MSNs in PD pathogenesis. LRFN2 is predicted to play a role in modulating chemical synaptic transmission and regulating postsynaptic organization, and is known to be important for brain development, function, and cognition^62–65^. To illustrate the genetic association of this *LRFN2* locus, we mapped all of the variants within the testing set and stratified them by their ML-predicted impact (**Fig. 5c**). PD individuals are enriched for predicted high-impact rare noncoding variants with many singleton variants observed in only one individual in the cohort (**Fig. 5c**). Further supporting a role for LRFN2 in PD pathogenesis, we find a significant decrease in *LRFN2* expression across the neuropathological progression of PD (**Supplementary Fig. 12d**). To illustrate variants that might potentially mediate this effect, we highlight two rare noncoding variants within the *LRFN2* locus identified within the GP2 cohort. These variants fall within regulatory elements and are predicted to disrupt their respective TF motifs—SIX5 (chr6:40532582, C to T) and SMAD4 (chr6:40513146, T to C) — thereby, reducing chromatin accessibility at their associated enhancers, consistent with the observed decrease in *LRFN2* expression across the neuropathological progression of PD (**Fig. 5d**).

Beyond having statistical power to associate novel genes with PD, our rare variant association testing methods also enable statistical association of rare noncoding variants within individual regulatory elements with PD (**Supplementary Fig. 12e**). While many of these associations overlap gene-based testing units, some are only uncovered at the level of individual regulatory element associations (**Supplementary Table 8**). Moreover, the number of peak regions tested varied across the different cell types, with no clear correlation between the number of peaks tested and the number of significant hits uncovered (**Supplementary Fig. 12b**). Overall, the use of ML prioritization enables the discovery of rare noncoding variants statistically associated with PD genetic risk and our results suggest that this platform is well-positioned to identify additional novel risk signals in larger discovery and replication cohorts.

### Nomination of novel noncoding variants driving familial PD

Previous work has demonstrated that ∼70% of familial early-onset PD cases and 80% of familial late-onset PD cases do not have a known genetic cause^21^. These studies defined familial PD as a family history of PD in up to a third-degree relative. The ability of ML models to rapidly prioritize noncoding variants makes them well-suited for understanding these familial cases of PD of unknown genetic origin. To this end, we analyzed families with multiple first-, second-, or third-degree relatives with a clinical diagnosis of PD but no coding alteration in a known PD risk gene. We performed genome-wide segregation analysis to identify all variants that tracked with disease status and focused our downstream analyses on two families where no plausible coding driver of PD could be identified.

In the first family, all three brothers developed PD by age 60, although no information was available about their parents (**Fig. 6a**). Due to the high genetic relatedness of the brothers, 7,283 variants segregated with the disease (**Fig. 6b**). Applying our ML models, we prioritized 102 unique noncoding variants within chromatin accessibility peaks. Of these, the variant with the strongest predicted impact resided within one of the promoters of the *PAQR5* gene and was predicted to exert its effects in microglia (**Fig. 6c–d**). Within the brain, *PAQR5* is expressed in microglia, perivascular macrophages, and capillary endothelial cells (**Fig. 6e**), and this rare variant is predicted to decrease the binding of an SPI1 transcription factor (**Fig. 6f**). *PAQR5* encodes a progestin and adiponectin receptor family member, and adiponectin has been shown to have a neuroprotective role in cellular and mouse models of ɑ-synucleinopathies^66^. Although this variant is an intriguing and plausible driver of disease, additional, high-impact noncoding variants were also predicted in this family, necessitating future functional genomics-based studies to validate these and identify the true causal variant (**Supplementary Table 9**).

**Figure 6.**
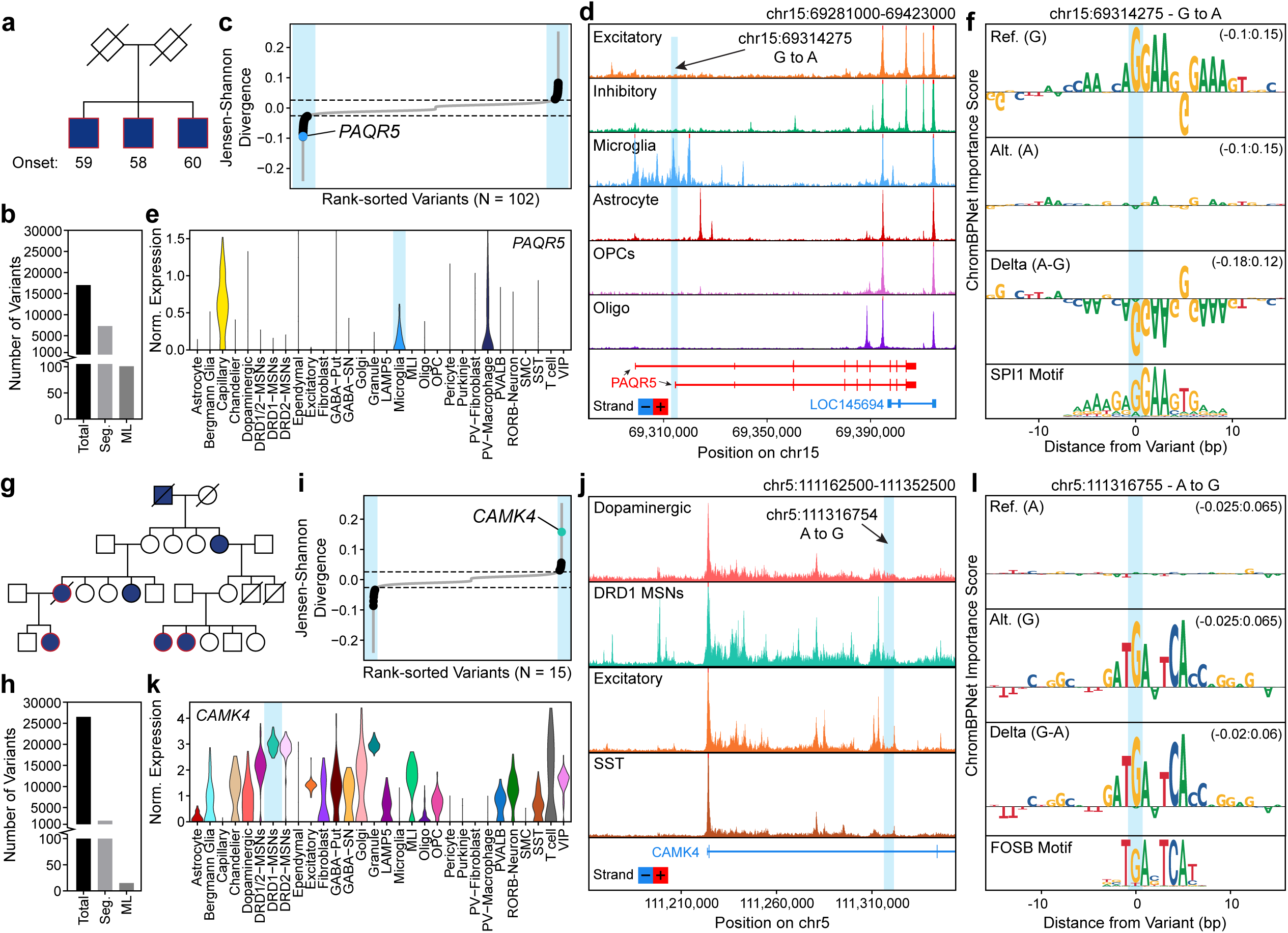
Rare noncoding variant prioritization identifies putative driver variants within PD-affected families. **a,g.** Pedigree diagram for (**a**) Family 1 and (**g**) Family 2 with affected individuals indicated by filled shapes. Squares represent biological males, circles represent biological females, and diamonds represent individuals for which additional information was not available. Age at onset is shown where available. **b,h.** Bar plot showing the total number of variants detected in (**b**) Family 1 and (**h**) Family 2, with subsets indicating rare variants that are shared among affected individuals (Seg.) and those predicted to have effects based on our ML models (ML). **c,i.** Rank-sorted ML-predicted effect scores for variants shared among affected individuals in familial PD cases. Prioritized effect variants found in (**c**) Family 1 and (**i**) Family 2 are highlighted with black points. The colored point in each plot indicates the prioritized effect variant – (**c**) chr15:69314275 - G to A in microglia in the *PAQR5* locus or (**i**) chr5:111316755 – A to G in DRD1 MSNs in the *CAMK4* locus. **d,j.** Normalized pseudobulked chromatin accessibility tracks across selected cell types for the (**d**) *PAQR5* locus and (**j**) *CAMK4* locus. The genomic position of the predicted causal variant is indicated with a blue highlight. **e,k.** Violin plot showing CPM-normalized expression of (**e**) *PAQR5* and (**k**) *CAMK4* across cell types. **f,l.** Importance score plots for (**f**) the *PAQR5*-associated variant and (**l**) the CAMK4-associated variant. Reference, alternate, and delta signals are shown, with the corresponding (**f**) SPI1 motif and (**l**) FOSB motif displayed below. Importance score ranges for each plot are shown in parentheses.

We additionally analyzed genetic data from a second family consisting of 4 affected individuals of more distant relatedness (**Fig. 6g**). This pedigree enabled a more robust segregation analysis, resulting in 1,895 segregating variants (**Fig. 6h**). Applying our ML models to segregating variants within chromatin accessibility peaks, we found 15 unique noncoding variants predicted to be functional (**Fig. 6i**). Of those, a variant intronic to the *CAMK4* gene arose as the likely causal variant, with a predicted effect in DRD1 MSNs (**Fig. 6j**). *CAMK4* is highly expressed in MSNs (**Fig. 6k**) and encodes a calcium/calmodulin-dependent protein kinase implicated in transcriptional regulation in neurons^67^. The variant is predicted to increase FOS-family transcription factor binding, potentially creating a novel gene regulatory element, and likely increase the expression of *CAMK4* (**Fig. 6l**). Interestingly, gain-of-function coding mutations in *CAMK4* have been associated with a neurodevelopmental disorder characterized by hyperkinetic movement^68,69^. Though PD is a hypokinetic movement disorder, the delicate balance of signaling between DRD1 and DRD2 MSNs controls proper movement, and thus perturbation of this balance may lead to either hypokinetic or hyperkinetic movement^70,71^. As such, we posit that this noncoding variant alters the expression of *CAMK4* in DRD1 MSNs and may play a role in driving the PD symptoms observed in this family. As shown by these examples, ML models can be leveraged to rapidly prioritize noncoding variants in familial pedigrees with disease of unknown genetic origin, opening the door to identification of novel risk genes through downstream functional genomic analyses.

## Discussion

Our study provides a framework for prioritizing and functionally interpreting rare noncoding variants in complex diseases. By integrating large-scale single-nucleus multi-omic profiling, whole-genome sequencing, and machine learning-based prediction, we bridge a long-standing gap in connecting noncoding genetic variation to cell type-specific regulatory consequences. In PD, where both common variant GWAS signals and rare coding variants have left much of the heritability unexplained, this approach nominates candidate genetic drivers across relevant cell types that have been previously overlooked. These findings underscore the importance of considering noncoding variation, not as background noise, but as a critical component of disease risk architecture.

A major advantage of this study is its comprehensive scale and resolution. We identified disease-associated alterations in both neuronal and non-neuronal cell populations by profiling over 3.3 million individual cell nuclei spanning five distinct brain regions. Our analysis confirms the known selective vulnerability of a specific dopaminergic neuron subset while simultaneously uncovering extensive transcriptional changes in glia and other neuronal subtypes. Further, cell type-specific QTLs and machine learning models developed from this dataset offer a robust platform for statistically assessing and predicting the functional impact of regulatory variants at scale. Notably, our machine learning-enhanced approach for rare variant association identifies genes and biological pathways that align with well-established PD mechanisms, including synaptic plasticity, protein trafficking, and cellular energy metabolism, while also highlighting previously uncharacterized cellular processes and novel candidate genes for further investigation. Importantly, the utility of our framework extends beyond sporadic disease to familial cases of unknown genetic origin. By applying our models to whole-genome sequencing data from familial PD pedigrees, we successfully nominated candidate causal variants in affected individuals, demonstrating the broad applicability of our approach for variant prioritization across different genetic architectures of PD.

Nevertheless, several limitations to our study should be considered. For one, although our schema allows for prioritization of noncoding variants within known chromatin accessibility peaks, it may not capture variants that generate completely novel regulatory elements. Second, although statistical association and allelic imbalance analyses confirm the functional impact of noncoding variants, experimental proof of causality is still required to elucidate the underlying molecular mechanisms. Moreover, while we use large sample sizes from studies such as GP2 and the UK Biobank, statistical power for rare variant association is inherently limited. A larger discovery cohort with many more control individuals would undoubtedly increase the replication rate of our findings in rare variant association testing. Similarly, many genetic signals may be masked by aggregation of variants into gene-centric testing units. The increased number of significant sets per cell type identified in our peak-based rare variant association testing suggests that PD genetic signals may be more readily uncovered when testing is performed on smaller, functionally coherent genomic units; however, this approach will require larger sample sizes and improved functional genomics tools to robustly interpret findings at the peak level. Future sequencing studies with larger cohorts, deeper multi-omic profiling, and functional perturbation experiments will be needed to refine variant prioritization and mechanistic understanding.

The approach outlined in this study—embedding multi-omic atlases, machine learning models, and rare variant association testing —offers a blueprint for dissecting the genetic basis of other neurodegenerative diseases and complex traits in which rare noncoding variation is likely also to have underappreciated contributions. By expanding these frameworks to a broader range of tissues and disease states, we enable an increasing capacity to isolate novel heritability, refine cellular pathogenic mechanisms, and ultimately influence prognosis and therapeutic opportunities.

## Acknowledgments

We thank the members of the Corces lab and the GP2 consortium for critical review of this manuscript. We also thank Mylinh Bernardi and Felicia Miller of the Gladstone Genomics Core and Eric Chow of the UCSF Center for Advanced Technology for their assistance throughout the project This project was supported by the Global Parkinson’s Genetics Program (GP2; https://gp2.org). GP2 is funded by the Aligning Science Across Parkinson’s (ASAP) (https://ror.org/03zj4c476) initiative and implemented by The Michael J. Fox Foundation for Parkinson’s Research (MJFF) (https://ror.org/03arq3225). For a complete list of GP2 members see https://doi.org/10.5281/zenodo.7904831. We thank Jennifer Jou, Khine Lin, Ben Hitz, Diane Trout, and Christopher McGinnis for guidance and assistance on data upload to the IGVF portal.

The Genotype-Tissue Expression (GTEx) Project was supported by the Common Fund of the Office of the Director of the National Institutes of Health, and by NCI, NHGRI, NHLBI, NIDA, NIMH, and NINDS. The data used for the analyses described in this manuscript were obtained from the GTEx Portal on 03/30/2025.

## Author Contributions

SM, AWT, and MRC conceived of the project. SM and MRC compiled the figures and wrote the manuscript with help and input from all authors. AWT, SHC, AWJ, HHC, AJS, and CC performed all the multi-omic data generation. AWT generated all the WGS data. SHC and LK generated all Micro-C data, and analysis of the Micro-C data was performed by SM, with the assistance of SHC. SM conducted all analyses of the multi-omic data and led all computational analyses. WGS data processing was performed by SM, ZHF, and CB. The UK Biobank analysis was conducted by YZ under the supervision of MEB. ZHF performed segregation analysis and data collection for the familial Parkinson’s disease data. AK and SD provided protocols and expertise related to fluorescence-activated nuclei sorting. CES and JMS provided expertise related to interpretation of genetic findings. TGB and GES curated the frozen tissue specimens used with input from TJM and MRC. MA, GP, RC, EMV, JT, and CG collected clinical samples from families with Parkinson’s disease and/or coordinated sequencing and data handling of relevant WGS data.

The full GP2 author list is presented in **Supplementary Table 10**.

## Data Availability

Whole genome sequencing data used in the preparation of this article were obtained from the Global Parkinson’s Genetics Program (GP2; https://gp2.org). Specifically, we used Tier 2 data from GP2 release 8 (https://doi.org/10.5281/zenodo.13755496). GP2 data can be requested through AMP PD (https://amp-pd.org).

All raw and processed snMultiome data is publicly available for download via the Impact of Genomic Variation on Function (IGVF) Data Portal at https://data.igvf.org/.

Please see **Supplementary Table 11** for details on data download. This includes a mapping of identifiers between IGVF and GP2 to enable cross referencing of whole genome sequencing data available and snMultiome data.

Processed snRNA-seq data is additionally available to download and explore on CellxGene at https://cellxgene.cziscience.com/collections/16876983-d454-43db-a8ec-77fafca9bf38. Tracks generated from our snATAC-seq data can be viewed through a WashU Epigenome browser session (https://epigenomegateway.wustl.edu/browser/?genome=hg38&sessionFile=https://files.corces.gladstone.org/Publications/2026_Menon_PDMultiome/WashU_sessions/ATAC_CellSubtype_WashU_SessionFile.json). ChromBPNet models, snATAC-seq peak sets, snATAC-seq bigwig track files, Micro-C loop calls, and peak-to-gene mapping files are deposited on Zenodo under accession number 10.5281/zenodo.17173379. FASTQ files from the Micro-C data are available at SRA BioProject PRJNA1418141.

## Code Availability

Analyses were performed using publicly available software. Statistical analyses were performed using R (v.4.1.0, v.4.2.1) available from the R site at https://cran.r-project.org/ and Python (v 3.11). All code generated for this article, and the identifiers for all software programs and packages used, are available on GitHub repository, https://github.com/GP2code/PDMultiome and were given a persistent identifier via Zenodo 10.5281/zenodo.18562016.

## Methods

### Genome annotations

All Whole Genome Sequencing, snATAC-seq, and snRNA-seq data were aligned and annotated to the hg38 reference genome.

### Generation of Whole Genome Sequencing libraries

For 26 of the individuals in this study, WGS had already been performed and was publicly available from the Global Parkinson’s Genetics Program (GP2). For the remaining 75 individuals, we isolated genomic DNA from cerebellum tissue pieces (<25 mg) and WGS libraries were made by Psomagen (Rockville, Maryland, USA). Genomic DNA isolation was performed using the QIAGEN DNeasy Blood & Tissue Kit. Samples with sufficient concentrations were assessed for DNA Integrity Number using the Agilent TapeStation 4200 and Agilent Genomic DNA ScreenTape. All samples sent for library preparation had a DNA Integrity Number greater than 7. WGS libraries were constructed using the Illumina TruSeq DNA PCR Free (350) Kit, assessed for quality using the TapeStation D5000 Screen Tape (Agilent), and quantified using qPCR before sequencing.

### Sequencing

Whole genome sequencing libraries were sequenced using an Illumina NovaSeq 6000 (S4 flow cell) by Psomagen (Rockville, Maryland, USA), each sample at 30x targeted coverage. Multiome snATAC-seq libraries were sequenced on either an Illumina NovaSeq 6000 instrument (S4 flow cell) or an Illumina NovaSeq X instrument (10B flow cell or 25B flow cell) using paired-end 50 bp reads. Multiome snRNA-seq libraries were sequenced on either an Illumina NovaSeq 6000 system (S4 flow cell) or an Illumina NovaSeq X system (10B flow cell or 25B flow cell) using 28 bp for Read 1 and 90 bp for Read 2. Micro-C libraries were sequenced on an Illumina NovaSeq X system (25B flow cell) using 2 x 150 bp reads. Sequencing was performed at the UCSF Center for Advanced Technology.

### Human patient cohort, sample acquisition, and patient consent

We performed single-nucleus multi-omic profiling of a cohort of 101 individuals with 5 brain regions per individual. This cohort consisted of 80 individuals with PD and 21 healthy controls. Cases and controls were matched for age and sex (full details of the cohort are provided in **Supplementary Table 1**). Primary, postmortem brain samples were acquired through the Banner Sun Health Research Institute Brain and Body Donation Program (Sun City, Arizona, USA) with institutional review board-approved informed consent. Sample sizes of cases and controls were chosen to provide confidence to validate methodological considerations. Putamen (subthalamic nucleus ranging to red nucleus), cingulate gyrus (anterior cingulate gyrus at Brodmann area (BA) 24/25/32/33), middle temporal gyrus (BA21), and cerebellum (taken of the folia of the sagittal cerebellum pole) were all obtained as frozen chunks. Substantia nigra was obtained as frozen cryosections (2 sections of 50 μm thickness each). Some brain regions were provided as larger tissue chunks that required additional manual cutting prior to nuclear isolation.

### Multiome experimental design and sample pooling approach

Before starting the project, we implemented an experimental design to reduce batch effects and mitigate differences between samples that could be due to technical variability. Instead of running each brain sample individually on one channel of the 10x Genomics chip, we implemented a sample pooling approach and ran the pooled nuclei across multiple channels of the chip. First, we defined a batch as a collective sample in which nuclei were isolated on the same day and in the same manner, counted individually, and subsequently pooled together in equal proportions. This pool subsequently underwent eight individual ATAC-seq transposition reactions and was loaded onto the same Chromium Next GEM Chip J across all eight channels. Most batches consisted of 12 samples.

To randomly assign samples to a particular batch, we used custom code that balanced different cohort characteristics. For the 505 samples, we randomized based on brain region, whether the sample was from an individual with PD or a control, biological sex, and ensured that an individual was only included once within a batch to enable genotype-based demultiplexing. Our experimental design also took into consideration having each of the five brain regions represented within an individual batch, as well as a consistent balance between PD cases and controls across all batches.

### Pilot studies to optimize nuclear isolation conditions

Before scaling up single-nucleus Multiome experiments with all 505 samples, we first conducted pilot experiments to evaluate several parameters. These included both snATAC-seq (transcription start site enrichment) and snRNA-seq (number of unique molecular identifiers (UMIs) per nucleus, number of genes per nucleus, levels of mitochondrial transcripts, levels of ambient RNA) metrics of data quality. Since our prior nuclear isolation protocol^72^ yielded relatively high proportions of oligodendrocytes from frozen brain tissue, our optimizations also aimed to capture a better balance of all brain cell types (different types of neurons, astrocytes, microglia, vascular cells) for downstream analyses. Therefore, using middle temporal gyrus samples, we benchmarked four nuclear isolation protocols for input into the 10x Genomics Multiome protocol.

1. The nuclear isolation protocol from Grandi et al. *Nature Protocols* 2022 using Dounce homogenization, an Iodixanol gradient (25%, 30%, and 40% Iodixanol layers), and homogenization buffer containing 0.3% NP-40.
2. The nuclear isolation protocol from Grandi et al. *Nature Protocols* 2022 using Dounce homogenization, an Iodixanol gradient (25%, 30%, and 40% Iodixanol layers), and homogenization buffer containing 1% Triton X-100 and Kollidon VA-64.
3. A nuclear isolation protocol adopted from the labs of Steve McCarroll and Evan Macosko (<u>dx.doi.org/10.17504/protocols.io.bi62khge)</u>. We used an extraction buffer containing 1% Triton X-100 and Kollidon VA-64. The frozen brain tissue was homogenized via trituration with a pipette on ice.
4. The 10x Genomics Chromium Nuclei Isolation Kit. For this protocol, frozen brain tissue was dissociated in the commercially provided Lysis Buffer and homogenizing with a plastic pestle. Debris was removed from using the provided Debris Removal Solution and nuclei were subsequently washed before resuspending in Diluted Nuclei Buffer and proceeding with transposition.

After the isolation of nuclei from samples using each of the four protocols, nuclei were processed using the 10x Genomics Multiome protocol. snATAC-seq and snRNA-seq libraries were generated, run on an Agilent BioAnalyzer for quality control, and sequenced separately on the Illumina NovaSeq 6000 using an S4 flow cell (UCSF CAT core). Samples generated using pilot protocols 1 and 2, which closely matched the final nuclei isolation protocol, were included in the final atlas; pilot protocols 3 and 4 were excluded from downstream analyses. These pilot samples were run in single-plex (1 sample per channel) rather than using genotype-based demultiplexing like the remainder of the atlas.<u>Isolation of nuclei from frozen tissue chunks and</u> <u>frozen tissue sections</u>

Nuclei were isolated from frozen tissue chunks and frozen cryosections. Our isolation protocol is available for public use at protocols.io at <u>dx.doi.org/10.17504/protocols.io.kxygxmr34l8j/v2.</u> This protocol is based on the nuclear isolation protocol in the Omni-ATAC protocol^72^ which was optimized exclusively for performing ATAC-seq on frozen tissue. To improve multi-omic data quality, we adopted several notable optimization strategies from published protocols for snRNA-seq of frozen brain. These included high-volume washes, buffer modifications, and low-speed centrifugation of the final nuclei pool at 250 g to prevent mechanical damage to nuclei, prevent leakage of valuable nuclear RNA transcripts, and reduce ambient RNA signals in the resultant Gene Expression libraries.

Briefly, frozen tissue segments were Dounce homogenized in 2 mL glass Dounces (Kimble) on ice to create a nuclei suspension and nuclei subsequently purified using an Iodixanol gradient. After density gradient centrifugation, a purified nuclei band was taken at the interface between the 30–40% Iodixanol layers. Nuclei were then washed and counted using the Countess 3 FL automatic counter (Thermo Fisher Scientific, Invitrogen) with ethidium homodimer I (Invitrogen, Thermo Fisher Scientific catalog E1169). Nuclei were differentiated from debris via ethidium homodimer staining and signal in the RFP channel of the Countess 3 FL. We took the RFP+ count of nuclei from each individual sample. After counting, nuclei were pooled together based on the above-mentioned batching strategy and subjected to a final high-volume wash prior to centrifugation and resuspension as input to the 10x Genomics Multiome protocol. Any remaining nuclei for each individual brain sample not used in pooling/transposition were cryopreserved by centrifugation for 10 minutes at 500 g at 4°C and resuspended in Bambanker Freezing Media (Wako Chemicals USA, Fisher Scientific catalog NC9582225). The nuclei resuspended in Bambanker were transferred to 1.8 mL Nunc Biobanking cryogenic tubes (Thermo Fisher catalog #375418), slowly frozen at −80°C overnight and subsequently stored at −80°C.

This isolation protocol was used for the vast majority of atlas generation, with the exception of pilot samples that were ultimately included in the final atlas. See the previous section on pilot studies for additional details regarding included nuclei.

### Re-profiling frozen nuclei samples

We sought to obtain a minimum of 3,500 pass-filter nuclei for each of the 505 brain samples. To achieve this, many samples required re-profiling. For brain samples with leftover nuclei from the initial isolation but no leftover tissue, we repeated single-nucleus Multiome reactions using the cryopreserved nuclei in Bambanker freezing medium. First, nuclei in Bambanker were removed from the −80°C freezer and placed immediately on ice in 1.8 uL cryovials. Next, we added cold wash buffer containing 0.1% Tween-20 to the cryovials and thawed the nuclei on ice. We again employed the pooling approach, combining nuclei from individual samples into a single 15 mL Falcon tube. These pooled nuclei were washed in the 15 mL Falcon tube, centrifuged for 10 minutes at 250 g at 4°C, supernatant removed, and the pellet resuspended in 1X Diluted Nuclei Buffer containing 1% BSA. The nuclei then underwent transposition and the 10x Genomics Multiome protocol according to the standard protocol.

### Micro-C experimental design and sample pooling approach

To obtain sufficient nuclei for Micro-C, we utilized middle temporal gyrus tissue samples from 36 individuals with PD, specifically with neocortical Lewy body pathology, and 21 healthy controls. Because microglia and astrocyte nuclei represent a small minority of the total nuclei recovered by Dounce homogenization, we pooled nuclei isolated from dozens of donors prior to sorting. In this pooling strategy, PD and control individuals were kept separate, but we did not attempt to uncover differences between the Micro-C signal of these groups. After isolating nuclei in the same manner as previously mentioned, nuclei were cryopreserved for later pooling and nuclei sorting. After combining sufficient nuclei, we employed fluorescence-activated nuclei sorting to obtain the 4 major brain cell types: microglia, astrocytes, oligodendrocytes, and neurons. Due to the low percentage yield of microglia and astrocytes from this process, the samples were split into 2 batches based on disease state with ∼150M nuclei each and sorted simultaneously using two BD FACS Aria Fusion machines. In this pooling process, the batches were optimized to balance the number of nuclei per individual to prevent bias toward any one donor. Biological and technical replicates for each cell type were then processed via Dovetail’s Micro-C Kit.

### Micro-C processing and analysis

Raw sequencing reads were aligned to the genome using BWA MEM. For each individual technical replicate, aligned reads were paired, sorted, and filtered for PCR duplicates and invalid pairs using Pairtools (v1.1.3) and then converted into pair files. Pair files across technical and biological replicates were merged to create a single file per cell type. The merged pairs files were used to normalize and generate matrix files with Juicer (v3.0). Chromatin contact loops were called using Mustache (v1.2.0) at 1, 5, 10, 20, 50, and 100kb resolutions.

### Fluorescence-activated nuclei sorting

The following protocol was applied to both batches separately. All samples were pooled to 150M nuclei in a 50 mL Falcon tube and mixed thoroughly. This mixed pool was then evenly distributed in 1.5 ml aliquots across multiple 15 ml Falcon tubes, each of which was then filled to 15 ml for a high-volume wash to dilute the cryopreservative. All pellets were resuspended in the same 10 mL of blocking buffer, and the pool was diluted to a concentration of 3.33M nuclei/mL, then stained with antibodies against nuclear markers for microglia (IRF5 AlexFluor 488-conjugated; R&D Systems IC4508G; final dilution 1:200), oligodendrocytes (SOX10; Bio-Techne custom conjugation to AF647; final dilution 1:200), and neurons (NeuN PE-conjugated, EMD Millipore FCMAB317PE; final dilution 1:1000) for 90 minutes, followed by DAPI (final concentration 1 ug/mL) for 15 minutes. Prior to antibody staining, 100k nuclei were removed and stained with DAPI alone for drawing population gates before nuclei sorting. No marker was used for astrocytes. The pool was concentrated into two 5 mL tubes of 75M nuclei each to be simultaneously sorted on two BD FACSAria Fusion Flow Cytometers. Each cell type was then pooled and concentrated separately in either 5 mL Eppendorf tubes or 15 mL Falcon tubes, depending on their volume and then pelleted and resuspended as input to the Dovetail Micro-C Kit.

### snATAC-seq and snRNA-seq data preprocessing

Reads were mapped onto the human reference genome (GRCh38) using CellRanger ARC (v2.02) software to obtain an aligned BAM, fragment, and raw_feature_bc_matrix files for each sample (10x Genomics). We then used demuxlet (via the Demuxafy^73^ v3.0.0 Singularity image) to deconvolute pooled nuclei using genotypes obtained from the 30x WGS. We then removed barcodes that were called as a doublet in either modality as well as barcodes that were identified as ambiguous in both modalities or cases in which the ATAC and RNA-assigned ID disagreed. snATAC-seq data was preprocessed using ArchR with fragment files as input.

Doublets were removed using the “filterDoublets” function and low-quality cells with a TSSEnrichment ≤ 3.5 or nFrags < 1000 were removed. We then manually reviewed each sample and increased the nFrags threshold to account for differences in sequencing depth. We added a Tile Matrix using ArchR’s default settings, followed by ArchR’s iterative Latent Semantic Indexing (LSI) dimensionality reduction with the top 80,000 variable peaks and resolution = 0.4. Cells were projected in 2D space using uniform manifold approximation projection (UMAP). Doublets on the snRNA-seq modality were removed using scds^74^ (v1.13.1 via Demuxafy v3.0.0 Singularity image) applied to the raw counts matrix. The snRNA-seq data were preprocessed using Seurat (v5.2.1), and single nuclei with fewer than or equal to 100 unique features (nGenes) or with greater than or equal to 10% of counts derived from mitochondrial genes were removed. Following the initial quality control, gene expression data were log-normalized and scaled.

### Identification of clusters and cell types

Cell type annotation was performed using a hierarchical, brain region-aware strategy leveraging paired single-nucleus chromatin accessibility and gene expression data. Initial cell classification was carried out using the chromatin accessibility modality. To account for brain region-specific regulatory landscapes, data were first split by brain region (Putamen, Substantia Nigra, Cerebellum, Middle Temporal Gyrus, and Cingulate Gyrus). Within each region, dimensionality reduction was performed using iterative LSI implemented in ArchR, using the top 25,000 most variable peaks across two iterations. Clustering was conducted using the Louvain algorithm with a resolution of 0.2.

Broad cell type identities were assigned per-cluster based on gene activity scores derived from chromatin accessibility profiles. For each cluster, module scores were computed using curated marker gene sets representing canonical cell types, including:

Excitatory Neuron (*RBFOX3, SYT1, SNAP25, TUBB3, MAP2, SLC17A7*)
Inhibitory Neuron (*RBFOX3*, *SYT1*, *SNAP25*, *TUBB3*, *MAP2*, *GAD1*, *GAD2*)
Other Neuron (*RBFOX3*, *SYT1*, *SNAP25*, *TUBB3*, *MAP2 but not in above categories)*
Oligodendrocyte (*MBP*, *MOG*, *PLP1*, *MAG*, *MOBP*, *CLDN11*)
OPC (*PDGFRA*, *CSPG4*, *OLIG1*, *OLIG2*, *SOX10*)
Astrocyte (*GFAP*, *AQP4*, *SLC1A2*, *SLC1A3*, *ALDH1L1*, *APOE*)
Microglia/Immune (*TMEM119*, *P2RY12*, *CX3CR1*, *SALL1*, *TREM2, C1QA*)
Vascular/Endothelial (*PECAM1*, *CLDN5*, *SLC2A1*, *FLT1*, *VWF*, *ENG*)

Each cluster was assigned to the broad cell type corresponding to the module with the highest average score, ensuring unbiased label propagation across regions. Region-specific neuronal populations were further annotated using tailored marker gene sets. Medium spiny neuron (MSN) markers (*PPP1R1B*, *GPR88*, *DRD1*, *DRD2*, *TAC1*, *PENK*, *ADARB2*) were scored exclusively in the putamen, granule cell markers (*GRIN2C*, *GABRA6*, *ZIC1*, *NEUROD1*, *LHX9*) in the cerebellum, and dopaminergic neuron markers (*TH*, *SLC6A3*, *NR4A2*) in the substantia nigra.

To incorporate cells with gene expression data but lacking paired chromatin accessibility data, we performed nearest-neighbor label transfer in the gene expression-derived reduced-dimensional space. Each RNA-only cell was mapped to its closest annotated multimodal neighbor and assigned the corresponding snATAC-seq-derived cell type label, enabling consistent annotation across modalities.

To verify and refine cell identities, we implemented a two-stage reclustering strategy. First, broad cell type assignments were validated independently in both the ATAC and RNA modalities by re-embedding each dataset and confirming concordance between cluster membership and marker gene expression. Second, each major cell type was subclustered individually to resolve finer cellular heterogeneity. For neuronal and vascular/endothelial populations, subclustering and subtype annotation were performed in the RNA modality, with Harmony applied to correct for donor- and region-associated batch effects. This allowed for the identification of transcriptionally distinct subpopulations, including D1 and D2 MSNs, dopaminergic neurons, and vascular subtypes.

### Chromatin accessibility peak calls

ArchR’s “addGroupCoverages” function was used to create pseudobulk replicates for each cell type. We then used MACS2 integration with ArchR’s “addReproduciblePeakSet” function, setting a q-value threshold of 0.01, to create a merged peak set. This process merged peaks called for each replicate to generate a consensus peak set, ensuring reproducibility across pseudobulk replicates within each cell type. Peaks were annotated using the “addPeakAnnotations” function which associates the peaks with genomic features such as promoters, enhancers, and gene bodies.

### Differential gene expression and accessibility testing

To identify differentially expressed genes between PD and control samples while accounting for potential confounding factors, we implemented a covariate-aware differential expression analysis using edgeR (v4.4.0). First, we performed principal component analysis on normalized pseudobulk expression data for each cell type, retaining high-variance genes. We then calculated correlations between the top five principal components and key covariates (biological sex, postmortem interval, and age at death), identifying significant associations (|*r*| ≥ 0.4, p ≤ 0.05). Covariates significantly correlated with expression variation (Pearson’s r *p*-value < 0.05) were incorporated into a negative binomial generalized linear model for differential expression testing. We used trimmed mean of M values (TMM) normalization and estimated dispersion parameters with edgeR’s “estimateDisp” function. Differential expression was determined via likelihood ratio tests (glmLRT) comparing PD to control samples. This approach allowed us to systematically control for non-disease factors while increasing statistical power to detect disease-associated transcriptional changes across cell types.

To determine whether individual cell types exhibited a significantly elevated burden of disease-associated transcriptional changes, we performed a permutation-based enrichment analysis. For each cell type and then each combination of cell type and brain region, we recorded the number of genes identified as significantly differentially expressed (FDR < 0.05) in the covariate-aware differential expression analysis described above. We then generated a null distribution of DEG counts by randomly permuting Parkinson’s disease and control labels across samples 100,000 times, rerunning the full differential expression analysis at each iteration using the same design matrix and covariate structure. This approach preserves the sample size, covariate associations, and gene expression variance within each cell type, while disrupting any true relationship between disease status and gene expression. For each permutation, we recorded the number of genes meeting the FDR threshold. Empirical p-values were computed for each cell type as the proportion of permutations in which the number of DEGs was greater than or equal to the observed number. Cell types with empirical p-values < 0.05 were considered to exhibit a statistically significant burden of differential gene expression in Parkinson’s disease.

To assess whether transcriptional changes were accompanied by alterations in chromatin accessibility, we performed a targeted differential accessibility analysis restricted to regulatory elements linked to differentially expressed genes. For each cell type, differentially expressed genes were identified and ATAC-seq peaks assigned to these genes using the peak-to-gene mapping strategy described below were then tested for differential accessibility between cases and controls using a negative binomial generalized linear model implemented in DESeq2 (v1.40.0). Size factors were estimated using the positive-counts method, Cook’s distance outlier filtering was disabled, and gene-wise dispersion estimates were used when global dispersion trend fitting was unstable. False discovery rate correction was applied within each gene, yielding a gene-level adjusted p-value for each peak.

To identify transcriptional signatures shared across brain regions within individual cell types, we examined the overlap of significantly differentially expressed genes (FDR < 0.05) identified in our covariate-aware DE analysis across the five profiled brain regions. For each cell type, we determined the number of genes that were differentially expressed in at least two regions (i.e., overlapping DEGs). To assess whether this degree of cross-region overlap exceeded what would be expected by chance, we implemented a permutation-based significance analysis.

Specifically, for each cell type, we used the previously generated permuted differential expression results (n = 10,000 permutations) to extract, for each iteration, the set of significant DEGs per region. Within each permutation, we then computed the number of genes appearing in two or more regions for that cell type, generating a null distribution of shared DEGs expected under the null hypothesis. We additionally examined the overlap between region-specific DEGs and subcluster-level DEGs, using hypergeometric testing to assess the statistical significance of shared genes and performing pathway enrichment analysis on the overlapping sets to identify convergent biological themes.

### Pathway Enrichment Analysis

Pathway and gene ontology enrichment analyses were performed using the enrichR^75^ (v3.3) R package. For each gene set of interest, over-representation analysis was conducted against curated biological databases, including GO_Biological_Process_2025, GO_Cellular_Component_2025, GO_Molecular_Function_2025, WikiPathways_2024_Human, and Reactome_Pathways_2024. Enrichments were computed using Fisher’s exact test with correction for multiple hypothesis testing as implemented in enrichR. For each database, adjusted *p*-values and combined scores were used to rank pathways, and significantly enriched terms were identified based on an adjusted *p*-value threshold (e.g., <0.05) unless otherwise specified.

### Cell proportion analysis

To assess whether the relative abundance of specific cell types varied across stages of Parkinson’s disease progression, we performed a differential composition analysis using the propeller() function from the *speckle* R package^76^ (v0.0.3). For each sample, we calculated the proportion of nuclei assigned to each annotated cell type, based on cell-level classifications from single-nucleus multiome data. To visualize these results in the CING and MTG, we generated heatmaps based on log_2_ ratio of PD to control mean proportions, with significant associations outlined. For significant cell types, we further explored trends across LB stages using linear regression.

### Expression changes association with LB stages

To identify genes whose expression changes with disease progression, we performed a trajectory-based differential expression analysis on pseudobulked RNA-seq profiles within each cell type using edgeR (v4.4.0). For each cell type, raw counts were aggregated at the sample level and filtered to retain genes with nonzero counts. Counts were normalized using the trimmed mean of M-values (TMM) method, and gene-wise dispersions were estimated using a negative binomial generalized linear model framework. We constructed a design matrix that included disease group and Unified Lewy body (LB) stage as a continuous covariate to model progressive disease effects. For each gene, we fit a negative binomial GLM and tested for (i) an overall disease effect using a likelihood ratio test on the disease group coefficient and (ii) a trajectory effect using a likelihood ratio test on the Unified LB stage coefficient. P-values were adjusted for multiple testing using the Benjamini–Hochberg false discovery rate procedure, and genes with FDR < 0.05 were considered significant.

For genes exhibiting significant trajectory effects, we further characterized expression patterns across disease stages by computing the mean normalized log-CPM expression within each Unified LB stage and classifying genes based on their behavior across the stages (e.g., monotonic increase, monotonic decrease). These patterns were used for downstream visualization and descriptive analyses of disease-associated transcriptional dynamics across cell types.

### QTL preprocessing and mapping

To identify cell type-specific QTLs, we focused on cell types with robust coverage across donors, requiring at least 10 individuals with more than 50 cells of that type. Expression QTLs (eQTLs) were called using TensorQTL^45^ and chromatin accessibility QTLs (caQTLs) were called using RASQUAL^44^.

For eQTLs, we extracted cell type-specific gene expression data from our Seurat object, excluding individuals with fewer than 50 cells per cell type. For each cell type, we aggregated single-cell RNA counts into pseudobulk expression matrices by summing counts across all cells per individual. We filtered genes to retain only those with non-zero expression in more than half of the samples and normalized expression data using log2-transformed counts per million (CPM). We prepared phenotype files in BED format compatible with TensorQTL, incorporating gene coordinates from GENCODE v46 and mapping gene symbols to Ensembl gene IDs. To account for technical and biological variation, we included biological sex, postmortem interval (PMI), age at death, Unified Lewy Body Stage, and the first five principal components derived from whole-genome sequencing data as covariates. Additionally, we performed principal component analysis on the normalized expression data and tested models incorporating 10, 20, 30, 40, 50, 60, 70, or 80 expression principal components as covariates. For inhibitory neurons, we combined VIP, PVALB, SST, and LAMP5 subtypes to increase sample size. We performed cis-eQTL mapping using TensorQTL, testing associations between genetic variants and gene expression within a 1 Mb window (±500 kb from the transcription start site). We used phased genotype data that passed quality control filters, with variants filtered to a minor allele frequency threshold of 0.05. We performed permutation testing with 10,000 permutations per gene to establish empirical significance thresholds and generate adjusted *p*-values for identifying significant eGenes. We ran TensorQTL in cis mode with permutation output to identify the lead eQTL per gene, while accounting for multiple testing.

For caQTLs, we extracted cell type-specific reads from our ArchR analysis in BAM format for each individual cell type, excluding cells with fewer than 50 reads from the analysis. To create feature sets for testing, we utilized cell type-specific peaks and converted them to SAF format. We applied featureCounts (v2.2.4) in paired-end mode to quantify reads per feature set, retaining features present in at least 50 individuals for each cell type. To streamline RASQUAL input file preparation, we used rasqualTools (https://github.com/marialauradias/rasqualTree/tree/master/rasqualTools) to generate compatible input files. We applied RASQUAL to calculate allele-specific counts per donor for each variant site by generating cell type-specific VCF files (createASVCF.sh). Using offsets, we corrected for both library size and GC content of each peak. We included biological sex, PMI, age at death, Unified Lewy Body Stage, and the first 5 genotype principal components as covariates. We used RASQUAL to call caQTLs, running the analysis with random permutations five times per cell type to establish empirical null distributions. To determine statistical significance, we filtered variants with squared correlation between the tested variant and feature SNPs (Sq_corr_rSNP) > 0.99, calculated *p*-values from chi-square statistics, and converted log10 q-values to q-values. For each cell type, we averaged the *q*-values across all five permutation runs and used these permuted q-values as the null distribution. We identified significant caQTLs at 5% FDR by comparing the observed q-value distribution to the average permuted *q*-value distribution.

### Enhancer-Gene mapping

We used the activity-by-contact model^77^ (ABC, v1.1.2) to identify putative enhancer–promoter interactions for cell types found in our dataset. We used our cell type-specific Micro-C data for chromatin conformation contacts data for microglia, astrocytes, oligodendrocytes and neurons. For OPCs, we used the publicly available averaged Hi-C data. For microglia, astrocytes, and oligodendrocytes, we used publicly available H3K27ac ChIP-seq data from Nott et al. 2019^52^. Enhancer activity was represented by the pseudobulked cell type-specific chromatin accessibility signal from our dataset. We used the default threshold of ABC score for the appropriate datasets provided. For Peak2Gene links, peaks were then linked to their target genes using ArchR’s “addPeak2GeneLinks” function to predict the regulatory relationships between identified peaks and their target genes.

### ChromBPNet model training and variant effect prediction

ChromBPNet models were trained using pseudobulked aggregate chromatin accessibility data from each cell type with more than 500 nuclei profiled. Bias models for each cell type were first trained on the negative background set in order to identify and regress Tn5 sequence bias.

These bias models were then used to train bias-corrected TF models across 5 different chromosomal splits. Variants were scored using predicted signals from the “profile” head of the models. Chromatin accessibility disruption scores are calculated using the Jensen–Shannon Divergence (JSD) between the predicted profiles of the reference sequence and the alternate sequence. The reference and alternate sequence are regions from the hg38 reference genome centered at the variant of interest. To identify variants predicted to exert the regulatory effects within each cell type, we implemented an approach based on inflection point analysis of the predicted JSD. For each cell type, the distributions of positive and negative JSD values were analyzed to identify the “elbow” (for positive scores) and “knee” (for negative scores) points along the ranked score curve. These inflection points correspond to the maximum perpendicular distance from the line connecting the first and last points of the normalized score distribution, thereby capturing the transition between the magnitudes of the background and outlier effects. Variants exceeding the positive elbow threshold or falling below the negative knee threshold were classified as “effect” variants for that cell type.

To account for variability in the number of regulatory elements associated with each gene, the count of effect variants was normalized by the number of accessible chromatin peaks mapped to that gene using cell type-specific peak-to-gene linkages. This normalization allowed comparison of variant burden across genes independent of differences in overall regulatory annotation density. To assess whether specific genes harbored more effect variants than expected by chance, we performed a permutation-based enrichment analysis. For each cell type, variant-to-peak mappings were randomly shuffled 100,000 times (maintaining per-gene peak counts) to generate an empirical null distribution of effect-variant counts. Empirical *p*-values were then computed as the proportion of permutations yielding equal or greater counts than observed, and these were adjusted for multiple testing using the Benjamini–Hochberg false discovery rate (FDR). Genes passing an FDR threshold of 0.1 were considered significantly enriched for cell type-specific effect variants.

To quantify the cell-type specificity of ChromBPNet-predicted effect variants, we aggregated all variant–peak–gene mappings across cell types and assigned each variant to a gene using a hierarchical rule that prioritized target-gene annotations (e.g., ABC-, correlation-, or distance-based links) and defaulted to the nearest gene when these annotations were unavailable but variants were located within 25 kb of the TSS. Variant-level JSD effect scores were merged with cell type-specific positive and negative thresholds, and variants were classified as effect variants if their scores exceeded the corresponding threshold for that cell type. To avoid double-counting due to multiple peaks linking the same variant to the same gene, effect variants were collapsed to unique (gene, variant, cell type) triplets. For each gene, we tabulated the number of unique effect variants observed in each cell type and computed both the per-cell-type proportion of variants and two specificity metrics: (i) the top-share (the maximum cell-type proportion) and (ii) the tau index, defined as 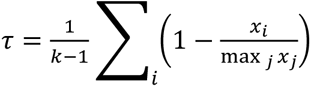, where *x*_*i*_ is the number of variants in cell type *i* and *k* is the number of cell types. To assess whether observed specificity exceeded chance, we constructed a permutation-based null model by shuffling cell type labels among effect variants within each gene while preserving the number of variants per gene; for each of 10000 permutations, we recomputed top-share and tau, yielding empirical null distributions from which 95th-percentile cutoffs were derived. Genes were considered cell-type-specific if either their observed top-share or τ exceeded these thresholds. We used pseudobulk RNA-seq profiles normalized to CPM; genes were evaluated only in cell types where CPM ≥ 1.

### Rare variant association testing

Rare variant association tests were performed using the optimal unified sequence kernel association test (SKAT-O) framework, as implemented in the “SKATBinary” function of the R SKAT package. Analyses were conducted separately for each cell type using both gene-based and peak-based variant groupings. For gene-based tests, variants were assigned to genes according to genomic annotations, whereas for peak-based tests, variants were grouped according to accessible chromatin regions identified within each cell type.

Prior to testing, variants were filtered to include only those with a minor allele frequency (MAF) < 0.01, based on allele frequencies from gnomAD (v4.1) for the discovery cohort (Global Parkinson’s Genetics Program; GP2) and from the UK Biobank (UKBB) for the replication cohort. Within each group, variants were further restricted to the top 1% of predicted functional effects according to machine learning-derived variant impact scores. Each variant was assigned a weight proportional to a combined function of its MAF and predicted functional effect, thereby up-weighting rarer variants with stronger predicted biological impact.

For each test, the binary phenotype (PD case versus control) was modeled while adjusting for biological sex and the first five genetic principal components (PCs 1–5) to account for ancestry. Association statistics were computed using the SKAT-O test, which adaptively combines burden and variance-component models to optimize power under heterogeneous effect directions. Associations found to be significant in an individual cell type in the GP2 cohort (prior to multiple hypothesis correction across cell types) were subsequently tested for replication in the UKBB cohort using the identical analytic framework and variant filtering criteria. For UKBB cohort selection, whole-genome sequencing (WGS) data from 490,276 UK Biobank participants were processed through a series of quality control and filtering steps to define PD cases and matched controls. After excluding individuals with sex-chromosome aneuploidy, excessive heterozygosity, or high missingness, 405,825 participants of White British ancestry remained. Participants with a PD diagnosis (N = 3,884) were compared to those without PD (N = 401,941). Among individuals with a PD diagnosis, we restricted our analysis to individuals with an age at onset greater than or equal to 60 years old (N = 3,337) and further retained only individuals without known PD driver variants (N = 3,071). Those PD driver variants were rs76763715 (*GBA1* N409S), rs75548401 (*GBA1* T369M), rs2230288 (*GBA1* E326K), rs34637584 (*LRRK2* G2019S), rs33939927 (*LRRK2* R1441S), rs104893877 (*SNCA* A53T), rs431905511 (*SNCA* G51D), rs104893875 (*SNCA* E46K), and rs104893878 (*SNCA* A30P). Kinship filtering (kinship coefficient > 0.0884) yielded a final set of 3,070 unrelated PD cases (1,941 males, 1,129 females). Among controls, individuals with dementia, other neurological disorders, or first-degree relatives with PD were excluded, and only those aged ≥65 years at baseline and without known PD driver variants (listed above) were retained (N = 55,247). Kinship filtering (kinship coefficient > 0.0884) yielded 54,526 unrelated individuals, from which a final set of 30,700 controls (19,410 males, 11,290 females) were 10:1 sex-matched to the above-mentioned PD cases for downstream analysis.

### Noncoding variant prioritization in familial PD cases

We analyzed WGS data from families containing more than three PD cases, which were part of the Global Parkinson’s Genetics Program (GP2) Data Release 8 (DOI: 10.5281/zenodo.13755496). We set genotypes to be missing after genotype quality control, defined as genotype quality >= 20, read depth >= 10, and heterozygous allele balance between 0.2 and 0.8, and retained high-quality variants with a call rate >= 0.95. Variant segregation analysis was performed using Slivar^78^ (v0.2.8) to identify variants consistent with dominant and recessive mode of inheritance within the families. We filtered the segregating variants against the Genome Aggregation Database (gnomAD; v4.0) genome dataset. For variants following a dominant mode of inheritance, we retained only those with a maximum credible population allele frequency <= 0.01. For variants consistent with a recessive mode of inheritance (homozygous or compound heterozygous), we retained those with a gnomAD homozygous variant count <=1. The segregating variants were then overlapped with peaks from each cell type and prioritized using the machine learning framework described above.

---

**Supplementary Figure 1.**
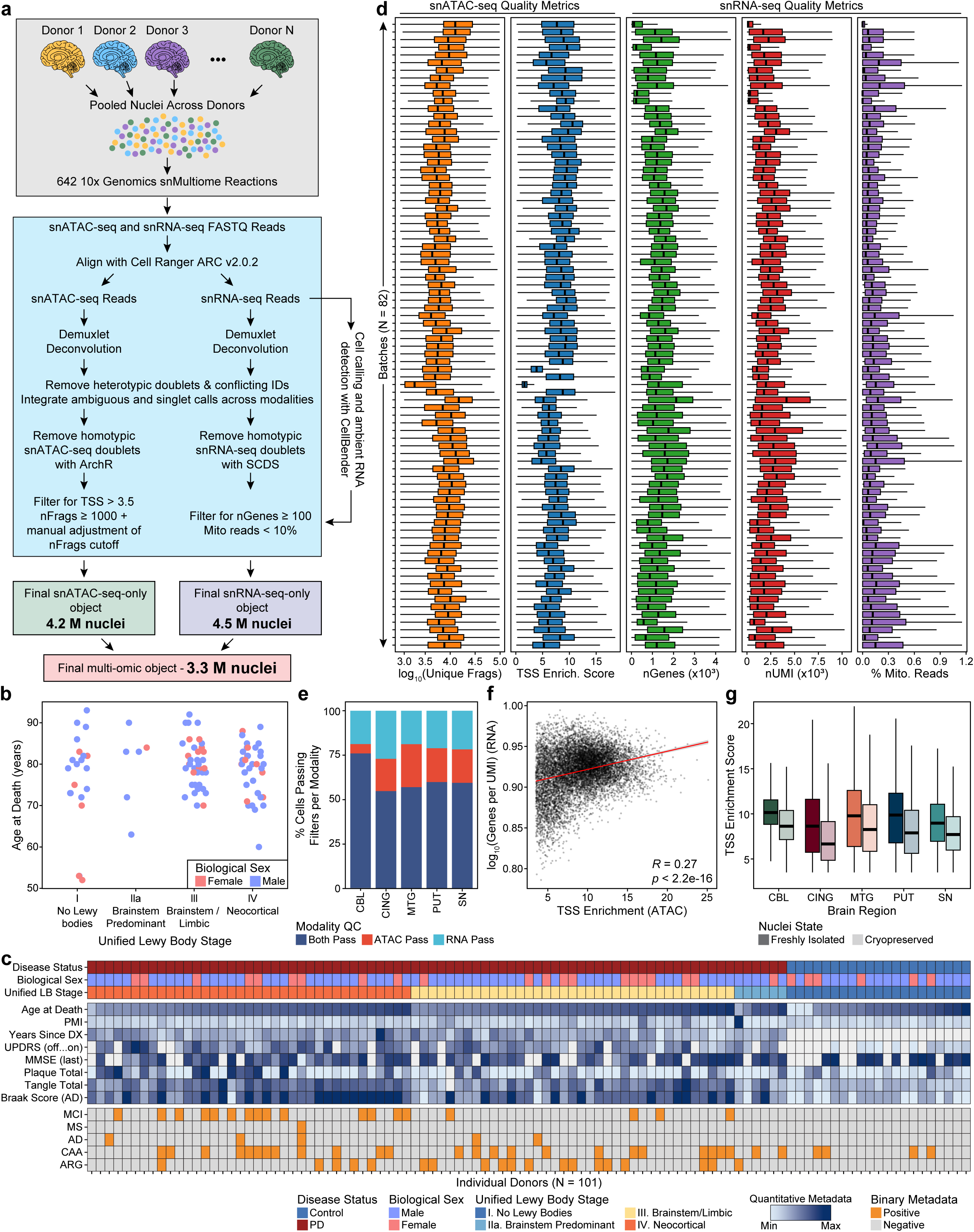
Multi-omic profiling of postmortem human brain yields high-quality and high-depth data across 101 donors. **a.** Schematic of workflow for snATAC-seq and snRNA-seq processing from pooled postmortem brain nuclei. Nuclei from multiple donors were pooled across 642 10x Genomics Multiome reactions. Reads were aligned, donors were demultiplexed, and doublets were removed separately per modality. Modality-specific quality control filters were then applied: snATAC-seq (TSS enrichment > 3.5, nFrags ≥ 1,000, with manual nFrags thresholding), and snRNA-seq (nGenes > 100, mitochondrial reads < 10%). Final datasets: snATAC-only (4.2M nuclei), snRNA-only (4.5 M nuclei), and multi-omic (3.3 M nuclei). **b.** Strip chart displaying donor distributions of age at death and biological sex across the Unified Staging System for Lewy Body Disorders. Each point is a donor, colored by biological sex (blue – male, pink – female). **c.** Matrix plot showing donor-level metadata. Columns show donors; rows include disease status, biological sex, Unified Lewy Body (LB) Stage, age at death, postmortem interval, years since diagnosis, clinical/neuropathology metrics, and quantitative region-specific pathology scores. **d.** Batch-level quality control for each modality. Box and whisker plots show (left to right within each modality): log_10_(unique fragments) (ATAC-seq; orange), TSS enrichment score (ATAC-seq; blue), number of detected genes (RNA; green), number of UMIs (RNA; red), and mitochondrial read percentage (RNA; purple). Each row corresponds to a processing batch. The center line denotes the median, the box spans the interquartile range (IQR; 25th–75th percentiles), and the whiskers extend to the most extreme data points within 1.5× the IQR. Outliers are not shown for cleaner visualization. **e.** Stacked bar plot displaying fraction of nuclei passing modality-specific quality control per brain region. **f.** Scatter plot of per-cell snATAC-seq TSS enrichment versus snRNA-seq genes per UMI for multi-omic nuclei; linear fit shown with Pearson correlation coefficient (r) and *p*-value from Pearson correlation test. **g.** Box and whisker plot comparing snATAC-seq TSS enrichment for nuclei freshly isolated from frozen tissue compared to nuclei isolated, cryopreserved, and then thawed across regions. The center line denotes the median, the box spans the interquartile range (IQR; 25th–75th percentiles), and the whiskers extend to the most extreme data points within 1.5× the IQR.

**Supplementary Figure 2.**
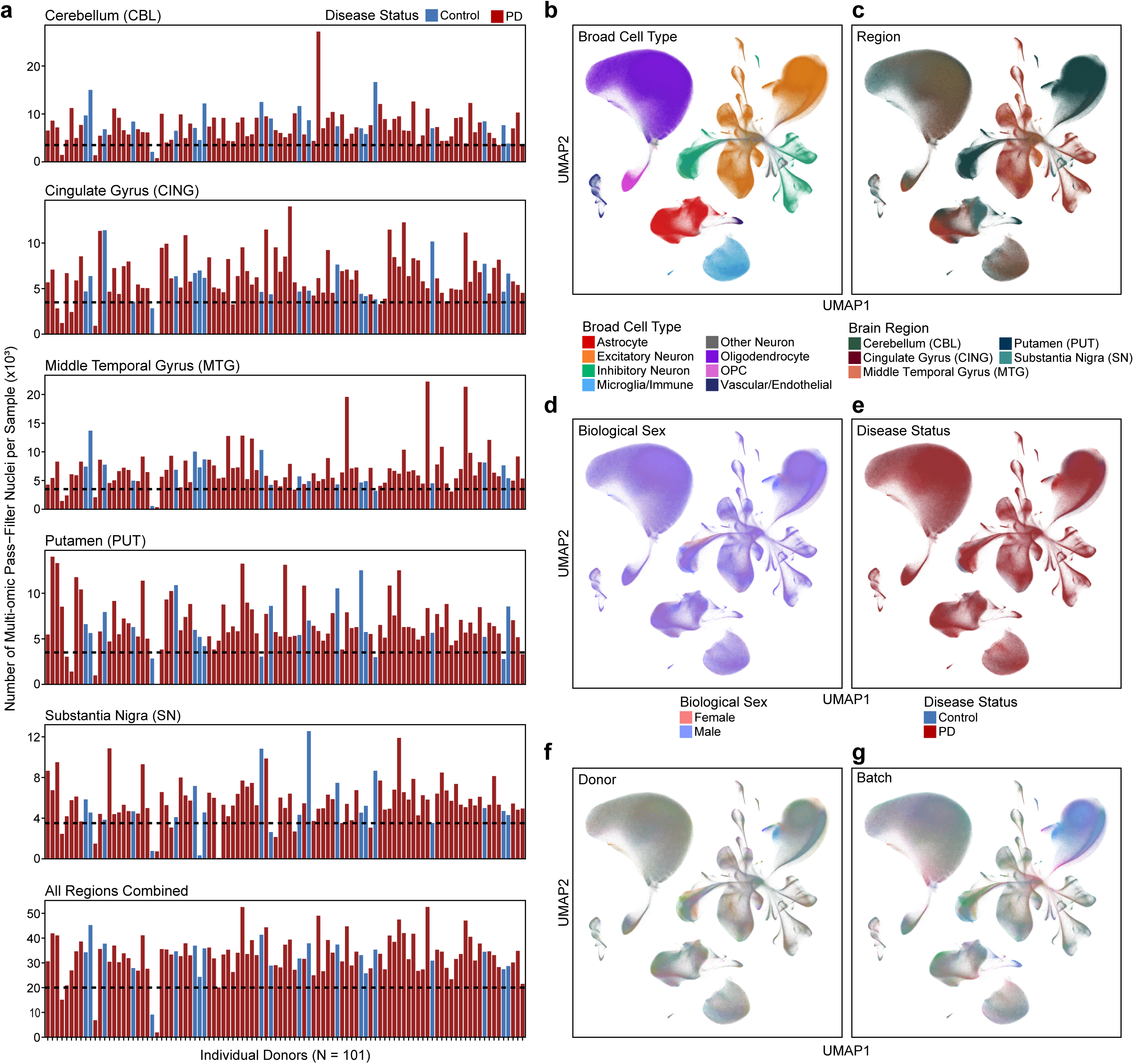
Sample multiplexing and genetic deconvolution enable uniform profiling across samples without observable batch effects across key covariates. **a.** Bar plot of the number of nuclei passing filtering thresholds per sample across each brain region. The bottom panel displays the combined totals across all regions for all 101 donors. Each bar represents one donor and is colored by disease status. **b–g.** UMAP embedding of nuclei colored by (**b**) broad cell type, (**c**) brain region of origin, (**d**) biological sex, (**e**) disease status, (**f**) donor identity, and (**g**) sequencing batch.

**Supplementary Figure 3.**
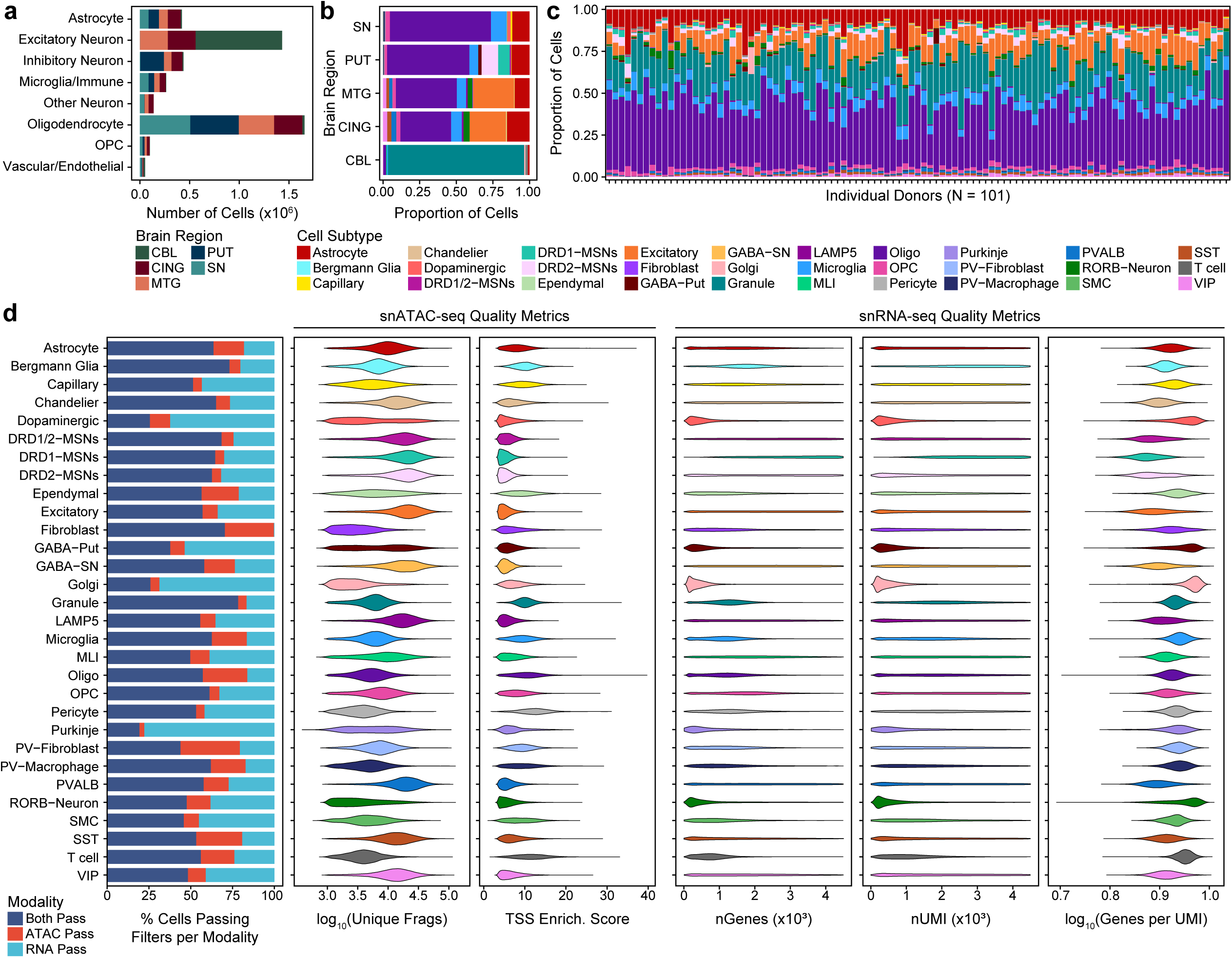
Cell type annotation identifies differences in brain regional composition and data quality across cell subtypes. **a.** Bar plot of the total number of nuclei per cell type across the dataset. Colors denote the brain region of origin. **b.** Bar plot of proportion of cell subtypes within each brain region. **c.** Stacked bar plot showing the proportion of cell subtypes contributed by each donor. Each column represents one donor. **d.** Quality control metrics displayed by cell subtype. In order from left to right: stacked bar plot showing the percent of nuclei passing modality-specific filters (ATAC-only pass, RNA-only pass, or passing both, violin plots of snATAC-seq metrics, including the number of fragments (nFrags) and TSS enrichment score, and violin plots of snRNA-seq metrics including number of detected genes (nGenes), number of UMIs (nUMI), and log_10_(genes per UMI).

**Supplementary Figure 4.**
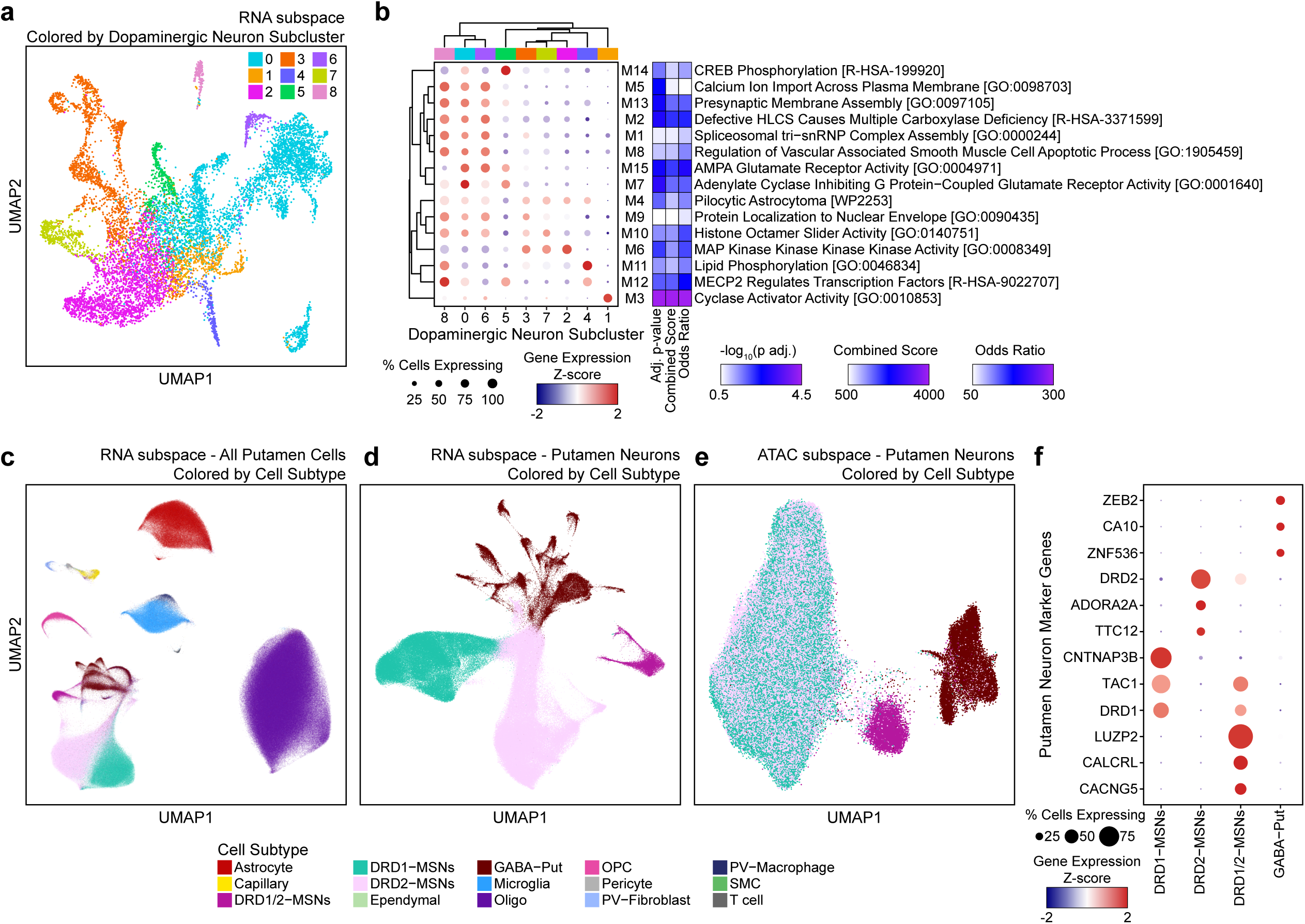
Subclustering of dopaminergic and putamen neuron populations uncovers subtypes of neurons with distinct molecular signatures. **a.** UMAP embedding of the RNA modality showing dopaminergic neurons, colored by subcluster identity. **b.** Dot plot showing subcluster representation across dopaminergic neuron modules. Point size indicates the percentage of cells expressing each module, and dot color encodes the average module eigengene value across clusters. Right panel: GO term pathway enrichment results for each module. **c.** UMAP embedding of (**c**) all putamen nuclei (RNA modality), colored by refined cell subtype, (**d**) putamen neurons (RNA modality), colored by refined neuronal subtype, and (**e**) putamen neurons (ATAC modality), colored by refined neuronal subtype. **f.** Dot plot of selected marker genes across putamen neuronal subtypes. Point size indicates the percentage of cells expressing each gene and point color indicates the z-scored average log-normalized expression level of each gene across all cells in that class.

**Supplementary Figure 5.**
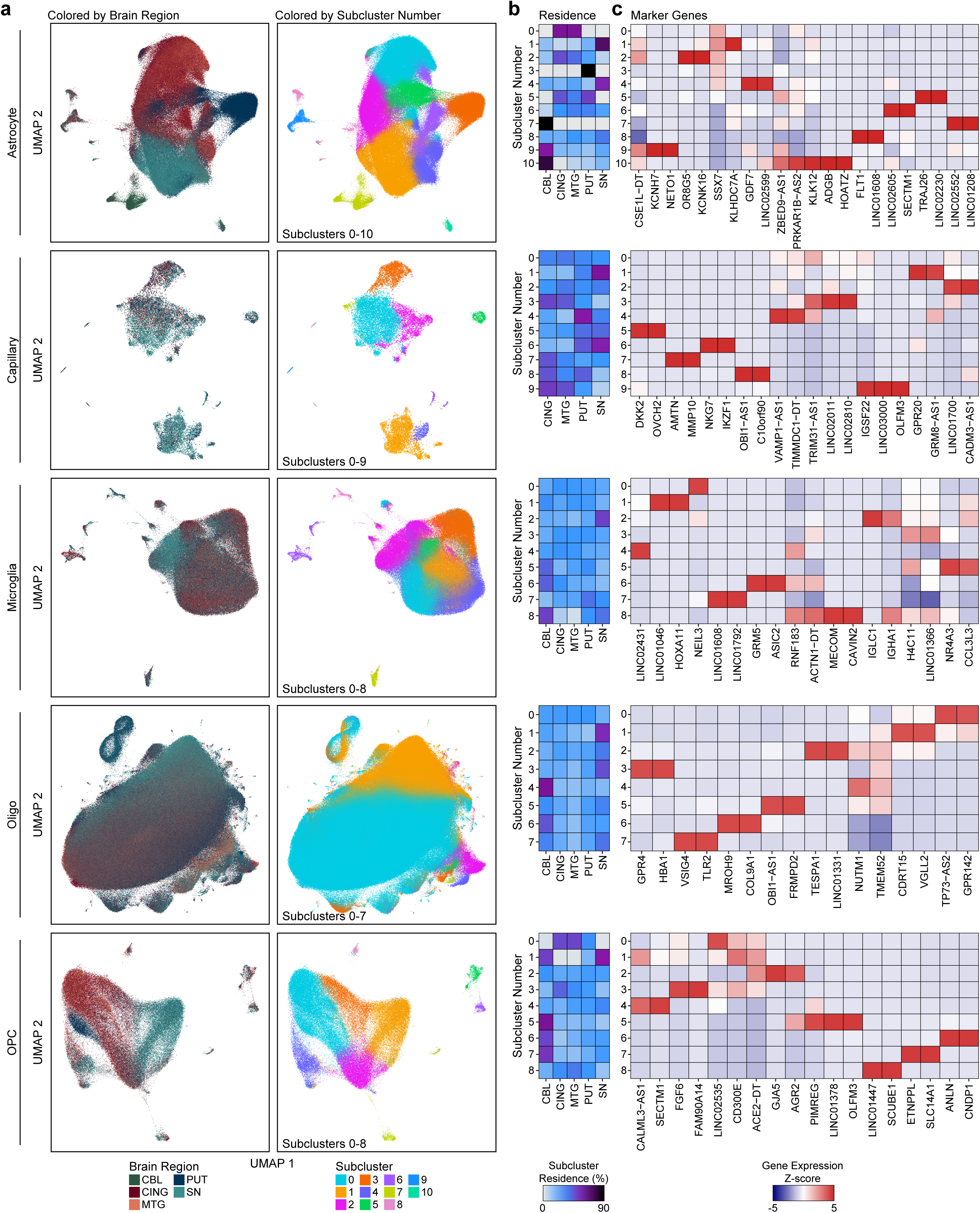
Subclustering and marker gene expression identify regional differences in non-neuronal cell populations. **a.** UMAP embeddings of astrocytes, capillary endothelial cells, microglia, oligodendrocytes, and oligodendrocyte precursor cells (OPCs) colored by (left) brain region and (right) subcluster identity. **b.** Cluster residence heatmaps showing the proportion of cells in each subcluster originating from each brain region, colored by percent of cells residing in each subcluster. **c.** Heatmaps of selected marker gene expression across subclusters for each glial population. Values are shown as z-score-normalized expression per gene.

**Supplementary Figure 6.**
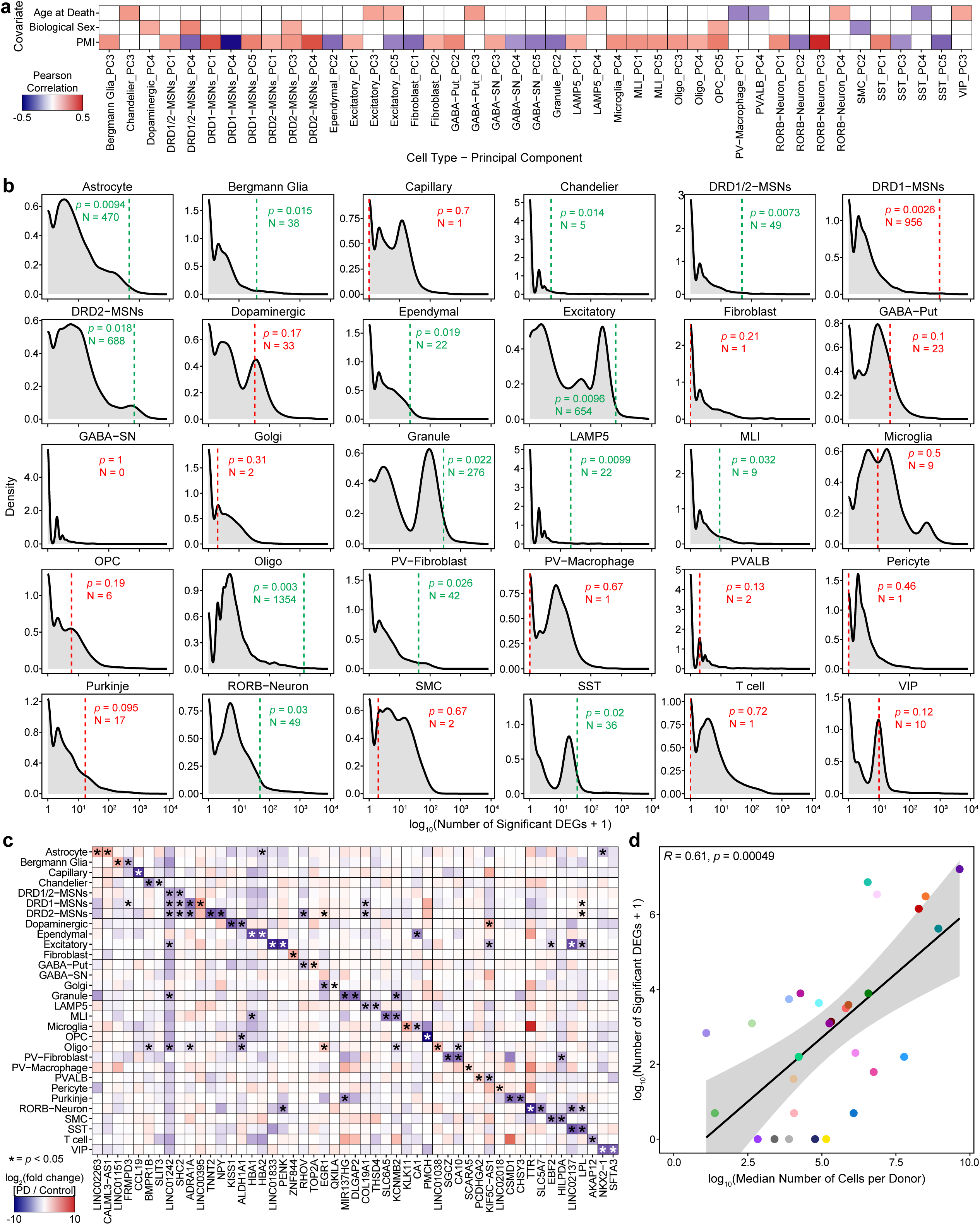
Differential gene expression analysis combined with permutation testing highlight cell types with statistically significant levels of molecular alteration between PD and control individuals. **a.** Heatmap displaying Pearson correlation coefficients between donor-level covariates (age at death, biological sex, postmortem interval (PMI)) and principal components (PC1–PC5) computed from pseudobulk expression profiles for each cell type. Values are shown per cell type/principal component. Principal components without significant correlation with covariates are excluded from the plot. **b.** Density plots showing the distribution of the number of significant differentially expressed genes (DEGs; log_10_-transformed) across 10,000 permutations of case/control label shuffling for each cell type. Vertical dashed lines indicate the observed number of DEGs in the real dataset. Reported *p*-values represent the proportion of permutations with equal or greater numbers of DEGs than observed; N indicates the number of donors for that cell type. Labels in green indicate a significant number of observed DEGs (*p* < 0.05); all other labels in red. **c.** Heatmap comparing log_2_(fold change) values of the top 2 DEGs by fold change across cell types. Rows represent cell types and columns represent genes. Color scale reflects the log_2_(fold change) of PD compared to Control. Asterisks indicate genes that are significant (FDR < 0.05) in the corresponding cell type based on a likelihood ratio test. **d.** Scatterplot comparing the median number of cells per donor and the number of DEGs identified as significant across the 29 tested cell types. Point color represents cell type as used throughout the manuscript. Fit line represents linear regression with the 95% confidence interval shown in gray.

**Supplementary Figure 7.**
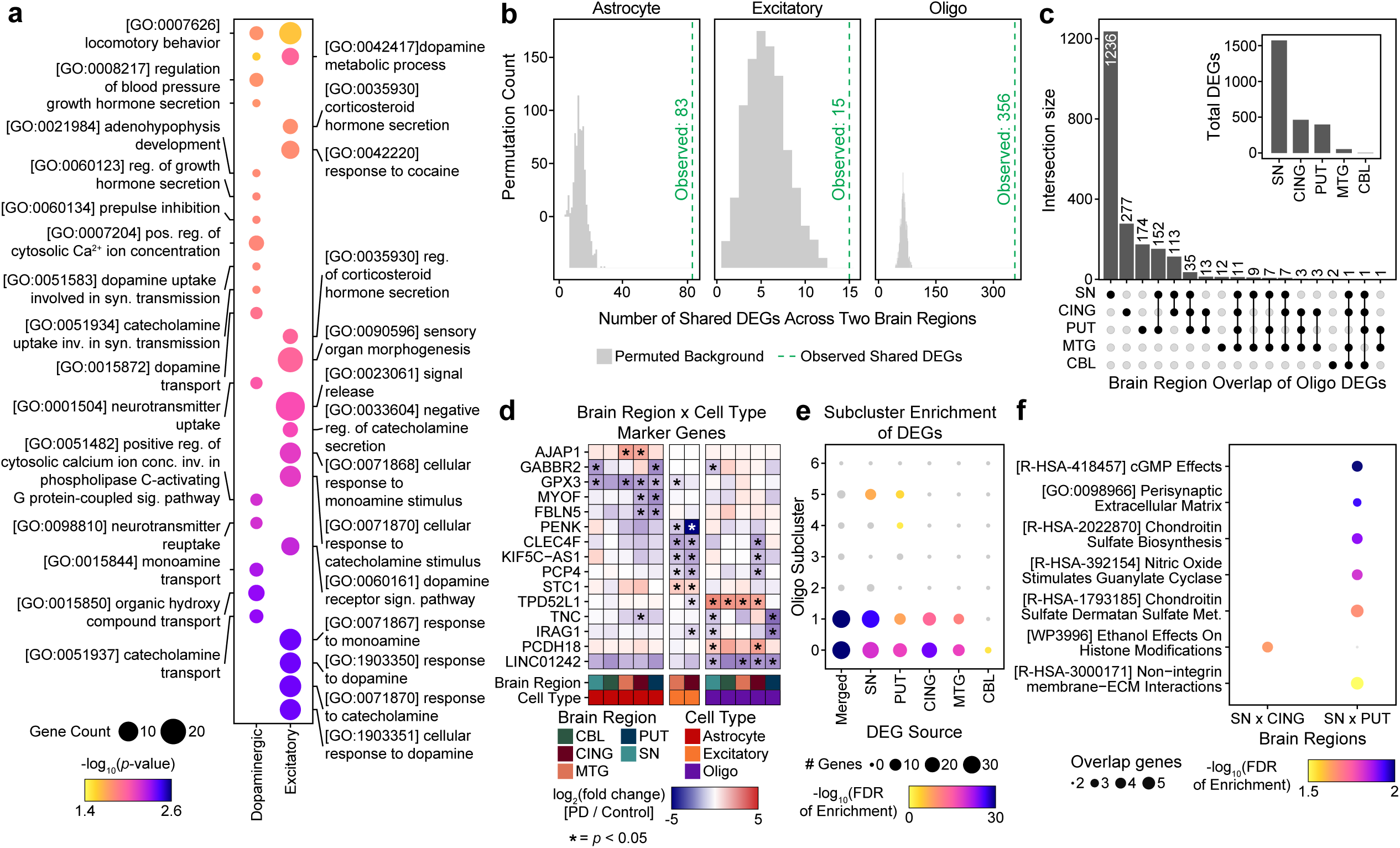
Differential gene expression analysis uncovers shared molecular signatures across brain regions. **a.** Dot plot of pathway enrichment of DEGs in Dopaminergic Neurons (left) and Excitatory Neurons (right). Point size represents the gene count per pathway; color represents the – log_10_(*p*-value) of enrichment. **b.** Histogram displaying permutation-based background distributions for the number of shared DEGs between pairs of brain regions in cell types with DEGs observed in multiple regions (Astrocytes, Excitatory Neurons, Oligodendrocytes). Dashed lines indicate the observed number of shared DEGs. **c.** UpSet plot showing overlap of DEGs across brain regions for Oligodendrocytes. Intersection sizes are shown on the left; the inset bar plot displays the total number of DEGs identified in each region. **d** Heatmap of selected marker genes displaying region-specific differential expression across cell types. Colors represent log_2_(fold change) of PD compared to Control; asterisks indicate significance (FDR < 0.05) based on a likelihood ratio test. **e.** Dot plot of enrichment of the overlap of region-specific DEGs within Oligodendrocyte subcluster DEGs. Point size represents the number of overlapping DEGs from each region and each subcluster, and color corresponds to the –log_10_ (FDR) of the enrichment of the overlap of genes within the two groups. Gray color indicates not significant. **f.** Dot plot of pathway enrichment of DEGs shared between brain region pairs in Oligodendrocytes. Point size represents the number of overlapping genes; color indicates –log_10_ (FDR) of the enrichment of overlapped genes within the given pathway. Gray color indicates not significant.

**Supplementary Figure 8.**
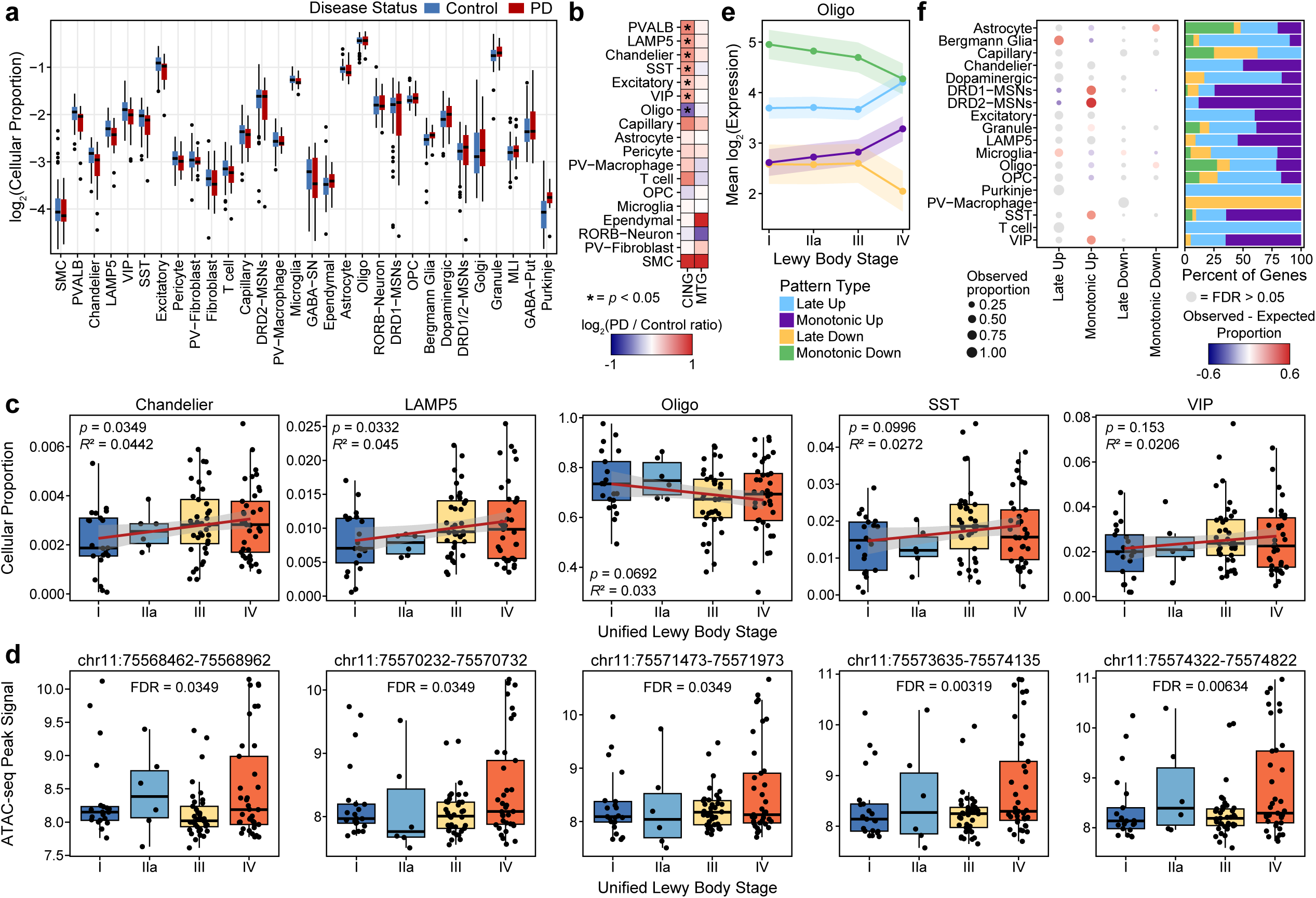
Cell type proportions and gene expression show distinct patterns across the trajectory of Lewy body accumulation in PD. **a.** Box and whisker plot showing log-transformed cell type proportions in control and PD donors. Each column represents a different cell type. Outlier donors are indicated as points. For box and whisker plots (**a,c,d**), the center line denotes the median, the box spans the interquartile range (IQR; 25th–75th percentiles), and the whiskers extend to the most extreme data points within 1.5× the IQR. **b.** Heatmap of the log_2_ (PD / Control) ratios of cell type proportions across the brain regions containing cell types that showed differential abundance in Figure 2c. **c.** Box and whisker plots showing cell type proportions across stages of the Unified Staging System for Lewy Body Disorders for interneuron and glial populations with significant difference in observed cell type proportions between PD and Control individuals (Chandelier, LAMP5, Oligodendrocytes, SST, VIP). Linear trend lines are shown with *p-*values and *R*² values from linear regression. **d.** Box and whisker plots of chromatin accessibility signal in Astrocytes at peaks mapping to *PRLR*, shown across stages of the Unified Staging System for Lewy Body Disorders. Each point represents a donor, and FDR values indicate the significance of stage association using likelihood ratio tests, which evaluate whether accessibility varies with Lewy body stage while adjusting for disease status. **e.** Plot of the mean log_2_ expression trajectories across stages of the Unified Staging System for Lewy Body Disorders for oligodendrocyte genes grouped by monotonic or late-stage expression patterns. Shaded regions represent the standard deviation. **f.** Left: Dot plot showing the proportion of genes in each cell type assigned to each temporal expression pattern category (late up, monotonic up, late down, monotonic down). Point size reflects the observed proportion; shading indicates deviation from the expected proportion. Right: Stacked bar plot showing the percentage of genes in each pattern category by cell type.

**Supplementary Figure 9.**
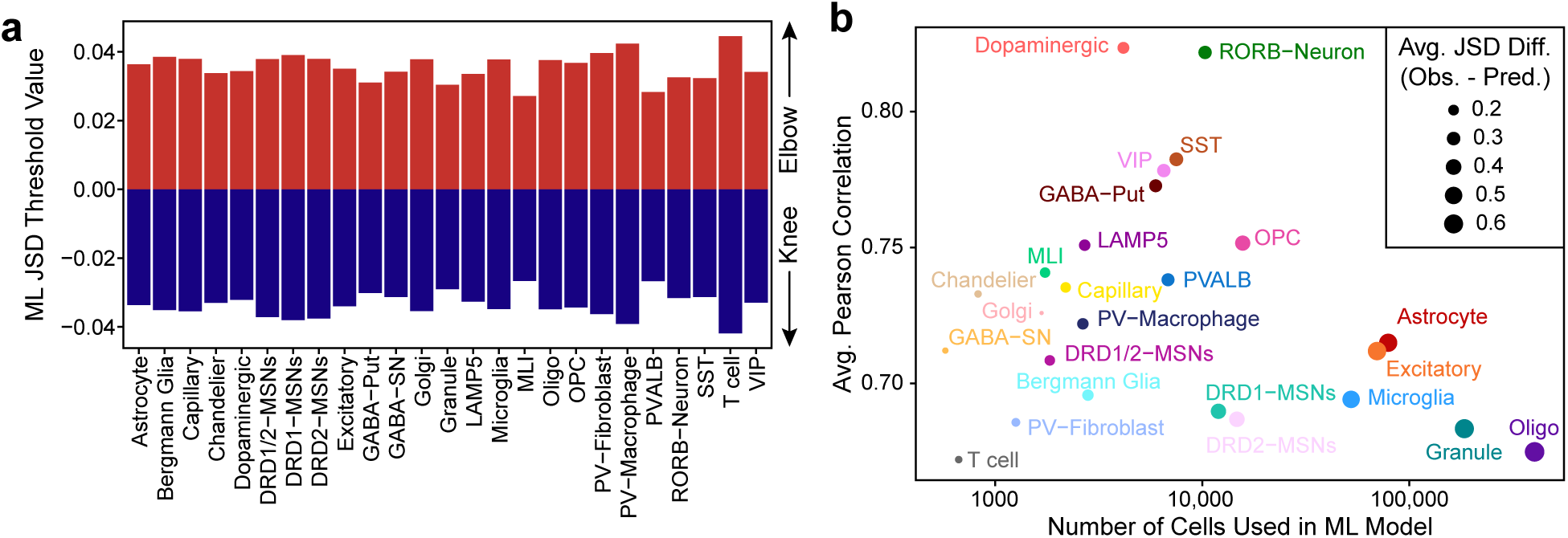
Cell type-specific ChromBPNet ML models learn chromatin accessibility patterns with comparable accuracy across cell types. **a.** Bar plot indicating variant effect size thresholds per cell type. Positive bars (red) indicate the threshold of positive effect (increased accessibility), and negative bars (blue) indicate the threshold of negative effect (decreased accessibility). **b.** Scatter plot of the average Pearson correlation of ML model predicted counts with observed counts across cross-validation folds plotted against the number of cells used for training, per cell type. Points are colored by cell type. Point size indicates the average median normalized Jensen–Shannon divergence (norm JSD) between observed and predicted accessibility profiles across regions. JSD values were normalized relative to a worst-case uniform prediction and a best-case perfect prediction (JSD = 0), yielding values between 0 and 1; larger points indicate closer agreement between predicted and observed profiles. This is averaged across 5 training folds.

**Supplementary Figure 10.**
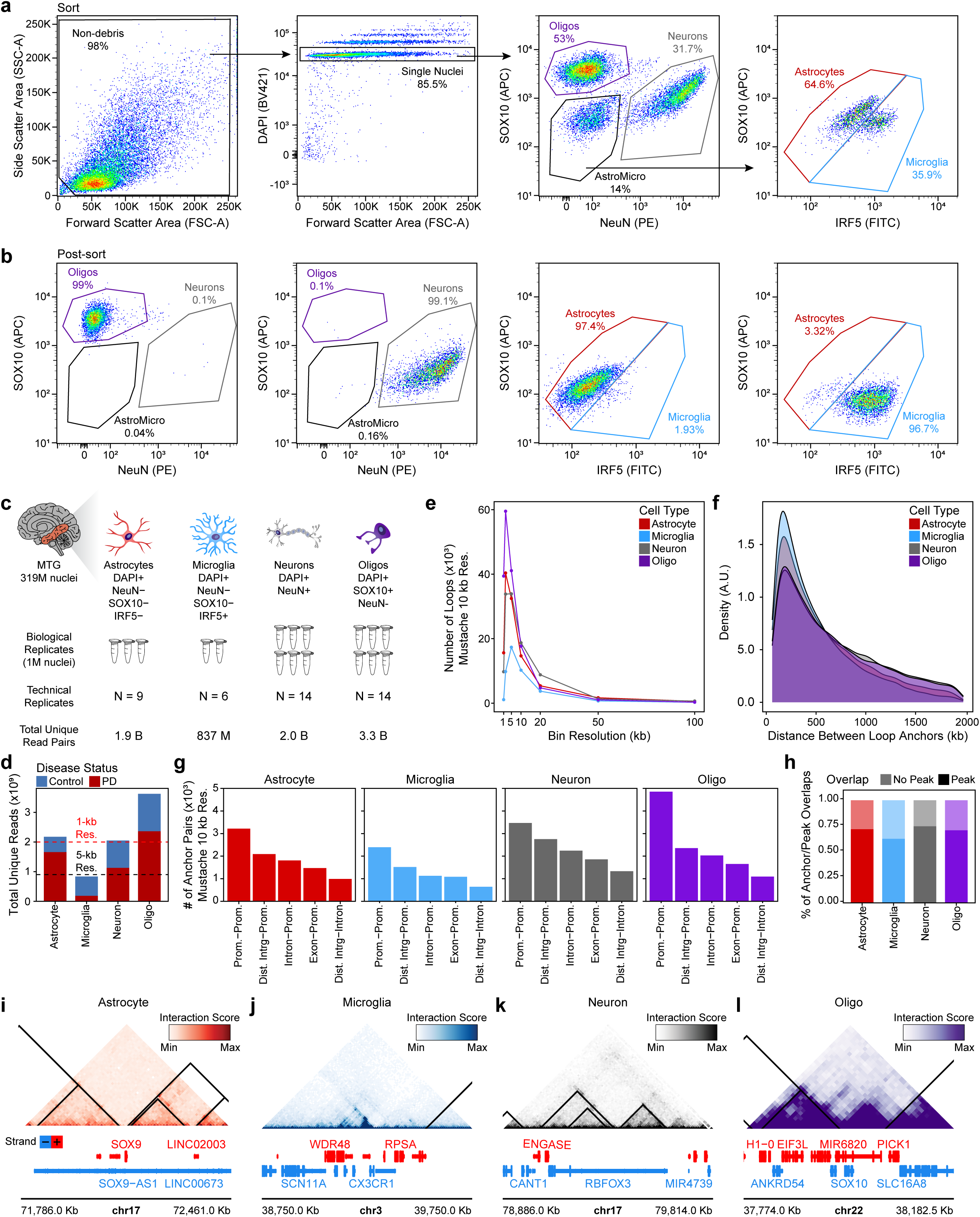
Cell type-specific Micro-C identifies variant-to-gene interactions. **a–b.** Representative fluorescence-activated nuclei sorting scatter plots of (**a**) the sorting strategy used and (**b**) the post-sort purity of the nuclei used for downstream Micro-C library generation. **c.** Schematic overview of Micro-C dataset generation. Cell type-specific nuclear markers used for sorting are indicated beneath each cartoon. **d.** Bar plot comparing total unique loops identified in Micro-C libraries generated from control samples and PD samples across astrocytes, microglia, neurons, and oligodendrocytes. Loop sizes are stratified by genomic span. **e.** Line plot showing the number of loops per 10-kb bin resolution, across astrocytes, microglia, neurons, and oligodendrocytes. **f.** Density plots of the distribution of genomic distances between loop anchors, across cell types. **g.** Bar plots of genomic annotations of loop anchor combinations per cell type. **h.** Stacked bar plot displaying the proportion of loop anchors overlapping accessible chromatin peaks in each cell type. **i–l.** Representative chromatin contact matrices for each cell type profiled, with contact intensities visualized as normalized Micro-C signal.

**Supplementary Figure 11.**
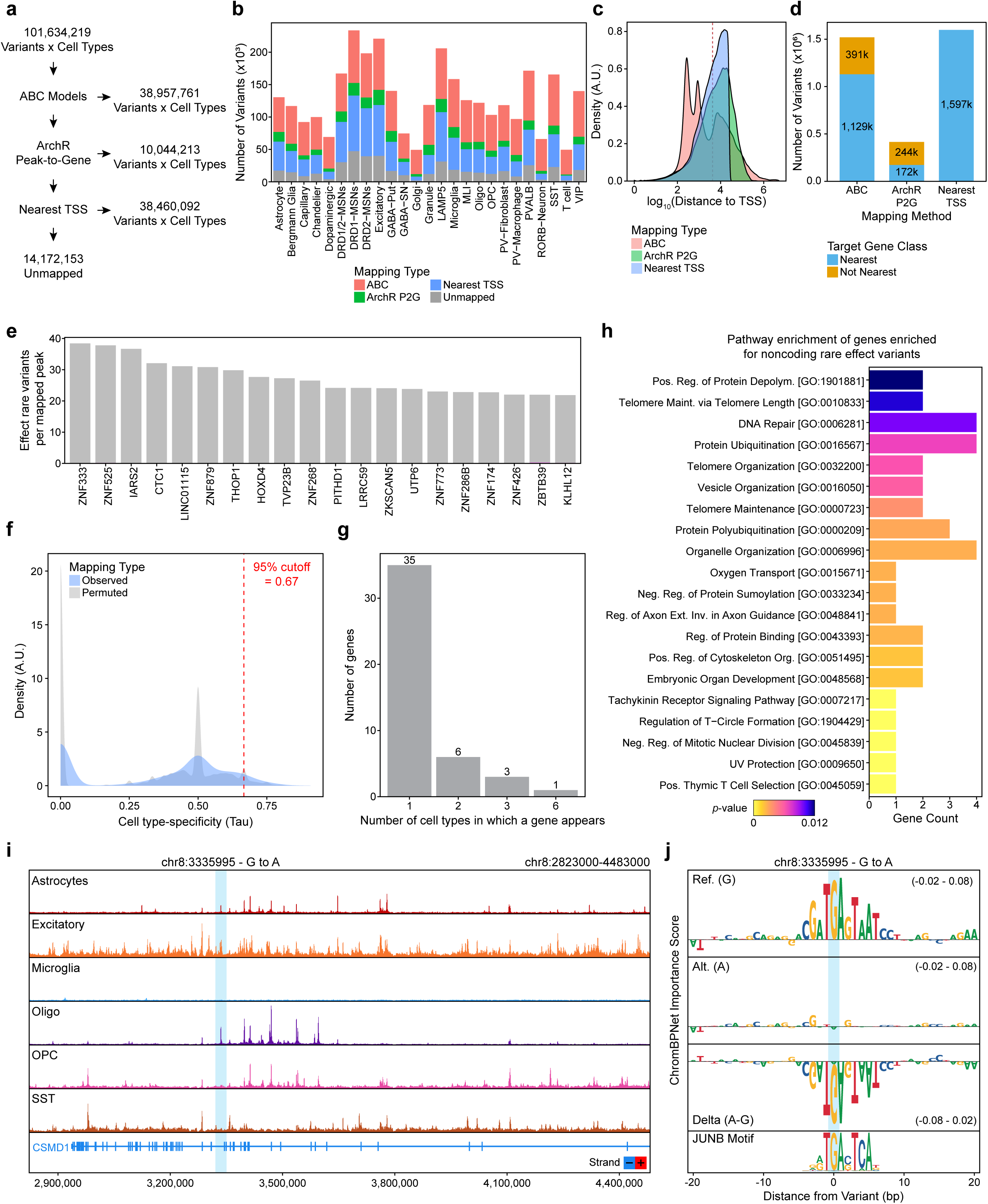
Multi-omic mapping links rare noncoding variants to precise, cell type-specific target genes. **a.** Overview of the variant-to-gene mapping workflow. Numbers shown refer to combinations of variants and cell types as some individual variants are assessed across multiple cell types. **b.** Stacked bar plot showing the number of rare variants (MAF < 0.01) identified as “hit variants” per cell type across the dataset. Stacked colors represent the mapping strategy used to assign the variant to its putative target gene. **c.** Density distributions of variant-to-TSS distances for each mapping strategy. The red vertical line denotes the median nearest-TSS distance. **d.** Stacked bar plot showing the number of variants linked to nearest versus non-nearest genes for each mapping method. **e.** Bar plot of the top 20 genes with the highest counts of predicted effect variants normalized by genomic area mapped to each gene per cell type. **f.** Distribution of cell type specificity (Tau) values for genes targeted by rare noncoding variant hits. Observed data (blue) and permuted background distributions (gray) are shown. The red dashed line marks the 95th percentile of the permuted distribution (Tau = 0.67), used as the significance cutoff. **g.** Histogram showing the number of cell types in which each gene is found to be enriched for rare noncoding variants. **h.** Pathway enrichment analysis of genes enriched for rare noncoding variant hits. Bars represent significant Gene Ontology terms, colored by *p*-value. **i.** Normalized pseudobulked chromatin accessibility tracks for the genomic region surrounding *CSMD1*. The area immediately surrounding variant chr8:3335995 – G to A is highlighted in light blue. **j.** Importance score plots for the variant shown in (**h**), including reference, alternate, and delta signal, with the associated JUNB TF motif shown below. Importance score ranges for each plot are shown in parentheses.

**Supplemental Figure 12.**
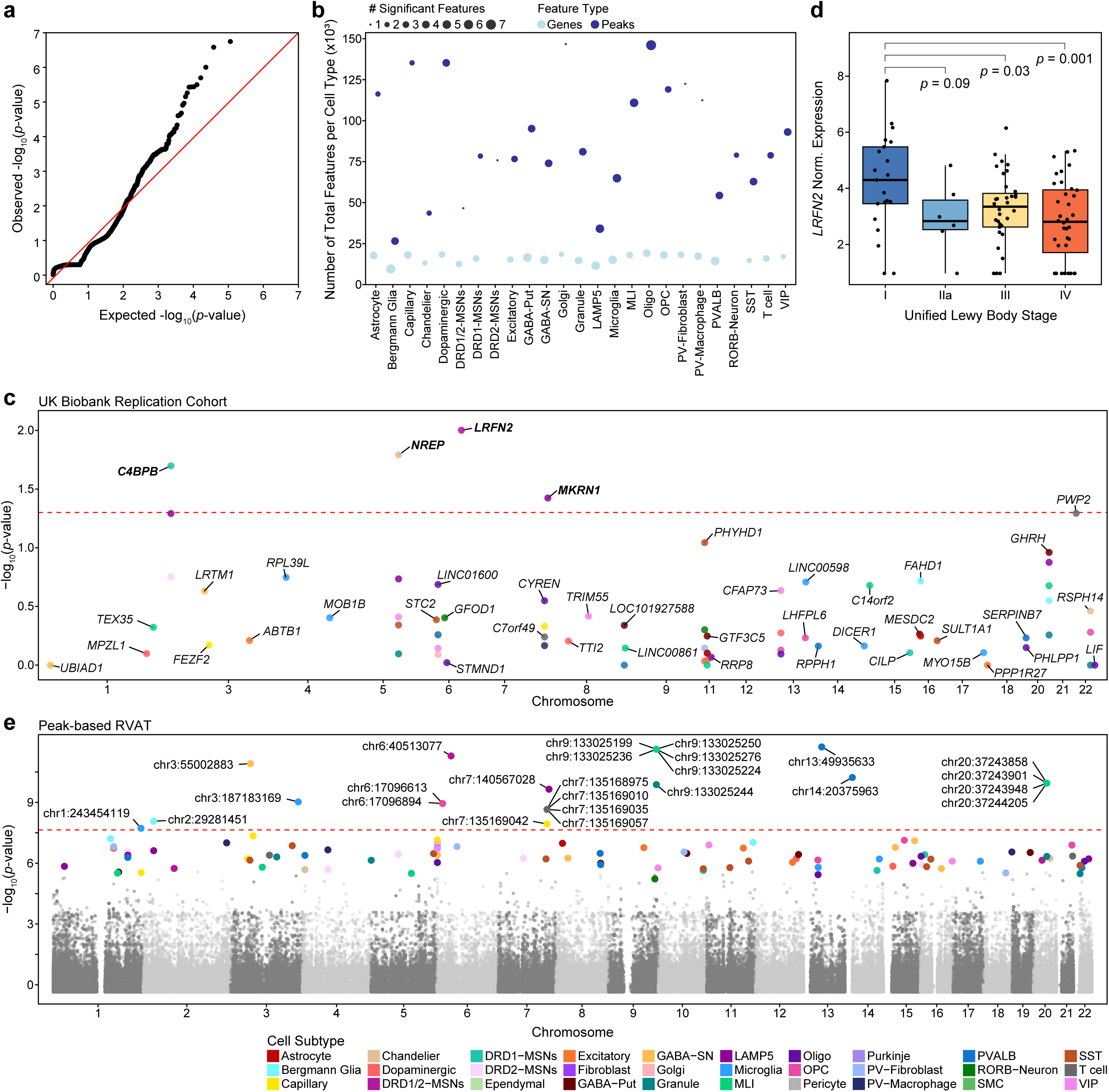
Rare variant association testing in PD identifies genes and putative regulatory elements enriched for noncoding variants. **a.** Quantile–quantile (QQ) plot of observed versus expected –log_10_(*p-*values) from a representative cell type-specific rare variant association test (shown for Oligodendrocytes). **b.** Dot plot of the number of significant rare variant-associated features per cell type. Point size reflects the number of significant features, and point color indicates whether the feature corresponds to a gene-set or a chromatin accessibility peak. **c.** Manhattan plot of rare variant association testing in the UK Biobank replication cohort of loci discovered in the GP2 cohort. The dashed red line denotes the replication significance threshold. Points are colored by cell type. **d.** Box and whisker plot showing log2(CPM+1) expression of *LRFN2* across stages of the Unified Staging System for Lewy Body Disorders. Pairwise stage differences were assessed using two-sample t-tests with pooled variance from a one-way ANOVA model; raw (unadjusted) *p*-values are shown. **e.** Manhattan plot of peak-based rare variant association testing with cell type-significant peaks colored by cell subtype. Labeled variants represent associations that are globally significant. Gray points indicate peaks that were not significant when tested in any individual cell type.

## References

1. Tolosa, E., Garrido, A., Scholz, S. W. & Poewe, W. Challenges in the diagnosis of Parkinson’s disease. Lancet Neurol. 20, 385–397 (2021).

2. Blauwendraat, C., Nalls, M. A. & Singleton, A. B. The genetic architecture of Parkinson’s disease. Lancet Neurol. 19, 170–178 (2020).

3. Nalls, M. A. et al. Large-scale meta-analysis of genome-wide association data identifies six new risk loci for Parkinson’s disease. Nat. Genet. 46, 989–993 (2014).

4. Kamath, T. et al. Single-cell genomic profiling of human dopamine neurons identifies a population that selectively degenerates in Parkinson’s disease. Nat. Neurosci. 25, 588–595 (2022).

5. Smajić, S. et al. Single-cell sequencing of human midbrain reveals glial activation and a Parkinson-specific neuronal state. Brain 145, 964–978 (2022).

6. Angelopoulou, E., Paudel, Y. N. & Piperi, C. Emerging role of S100B protein implication in Parkinson’s disease pathogenesis. Cell. Mol. Life Sci. 78, 1445–1453 (2021).

7. Martirosyan, A. et al. Unravelling cell type-specific responses to Parkinson’s Disease at single cell resolution. Mol. Neurodegener. 19, 7 (2024).

8. Gebremedhin, K. G. & Rademacher, D. J. Histone H3 Acetylation in the Postmortem Parkinson’s Disease Primary Motor Cortex. Neurosci. Lett. 627, 121–125 (2016).

9. Gordon, R. et al. Inflammasome inhibition prevents α-synuclein pathology and dopaminergic neurodegeneration in mice. Sci. Transl. Med. 10, eaah4066 (2018).

10. Nalls, M. A. et al. Identification of novel risk loci, causal insights, and heritable risk for Parkinson’s disease: a meta-analysis of genome-wide association studies. Lancet Neurol. 18, 1091–1102 (2019).

11. Chang, D. et al. A meta-analysis of genome-wide association studies identifies 17 new Parkinson’s disease risk loci. Nat. Genet. 49, 1511–1516 (2017).

12. Kim, J. J. et al. Multi-ancestry genome-wide association meta-analysis of Parkinson’s disease. Nat. Genet. 56, 27–36 (2024).

13. Maurano, M. T. et al. Systematic localization of common disease-associated variation in regulatory DNA. Science 337, 1190–1195 (2012).

14. Gaulton, K. J., Preissl, S. & Ren, B. Interpreting non-coding disease-associated human variants using single-cell epigenomics. Nat. Rev. Genet. 24, 516–534 (2023).

15. Kuksa, P. P. et al. Scalable approaches for functional analyses of whole-genome sequencing non-coding variants. Hum. Mol. Genet. 31, R62–R72 (2022).

16. Bloem, B. R., Okun, M. S. & Klein, C. Parkinson’s disease. The Lancet 397, 2284–2303 (2021).

17. Makarious, M. B. et al. Large-scale Rare Variant Burden Testing in Parkinson’s Disease Identifies Novel Associations with Genes Involved in Neuro-inflammation. 2022.11.08.22280168 Preprint at 10.1101/2022.11.08.22280168 (2022).

18. Bandres-Ciga, S., Diez-Fairen, M., Kim, J. J. & Singleton, A. B. Genetics of Parkinson’s disease: An introspection of its journey towards precision medicine. Neurobiol. Dis. 137, 104782 (2020).

19. Klein, C. & Westenberger, A. Genetics of Parkinson’s Disease. Cold Spring Harb. Perspect. Med. 2, a008888 (2012).

20. Lim, S.-Y. & Klein, C. Parkinson’s Disease is Predominantly a Genetic Disease. J. Park. Dis. 14, 467–482.

21. Towns, C. et al. Parkinson’s families project: a UK-wide study of early onset and familial Parkinson’s disease. NPJ Park. Dis. 10, 188 (2024).

22. Auton, A. et al. A global reference for human genetic variation. Nature 526, 68–74 (2015).

23. Hernandez, R. D. et al. Ultrarare variants drive substantial cis heritability of human gene expression. Nat. Genet. 51, 1349–1355 (2019).

24. Kelley, D. R., Snoek, J. & Rinn, J. L. Basset: learning the regulatory code of the accessible genome with deep convolutional neural networks. Genome Res. 26, 990–999 (2016).

25. Zhou, J. & Troyanskaya, O. G. Predicting effects of noncoding variants with deep learning–based sequence model. Nat. Methods 12, 931–934 (2015).

26. Ansari, M., Fischer, D. S. & Theis, F. J. Learning Tn5 Sequence Bias from ATAC-seq on Naked Chromatin. in Artificial Neural Networks and Machine Learning – ICANN 2020 (eds Farkaš, I., Masulli, P. & Wermter, S.) 105–114 (Springer International Publishing, Cham, 2020). doi:10.1007/978-3-030-61609-0_9.

27. Buenrostro, J. D., Giresi, P. G., Zaba, L. C., Chang, H. Y. & Greenleaf, W. J. Transposition of native chromatin for fast and sensitive epigenomic profiling of open chromatin, DNA-binding proteins and nucleosome position. Nat. Methods 10, 1213–1218 (2013).

28. Walker, L., Stefanis, L. & Attems, J. Clinical and neuropathological differences between Parkinson’s disease, Parkinson’s disease dementia and dementia with Lewy bodies – current issues and future directions. J. Neurochem. 150, 467–474 (2019).

29. Stolt, C. C. et al. The Sox9 transcription factor determines glial fate choice in the developing spinal cord. Genes Dev. 17, 1677–1689 (2003).

30. Smith, A. M. et al. The transcription factor PU.1 is critical for viability and function of human brain microglia. Glia 61, 929–942 (2013).

31. Furukawa, T., Morrow, E. M. & Cepko, C. L. Crx, a novel otx-like homeobox gene, shows photoreceptor-specific expression and regulates photoreceptor differentiation. Cell 91, 531–541 (1997).

32. Saunders, A. et al. Molecular Diversity and Specializations among the Cells of the Adult Mouse Brain. Cell 174, 1015–1030.e16 (2018).

33. Bergonzoni, G., Döring, J. & Biagioli, M. D1R- and D2R-Medium-Sized Spiny Neurons Diversity: Insights Into Striatal Vulnerability to Huntington’s Disease Mutation. Front. Cell. Neurosci. 15, 628010 (2021).

34. Buddhala, C. et al. Dopaminergic, serotonergic, and noradrenergic deficits in Parkinson disease. Ann. Clin. Transl. Neurol. 2, 949–959 (2015).

35. Dehestani, M. et al. Transcriptomic changes in oligodendrocytes and precursor cells associate with clinical outcomes of Parkinson’s disease. Mol. Brain 17, 56 (2024).

36. Bae, E.-J., Pérez-Acuña, D., Rhee, K. H. & Lee, S.-J. Changes in oligodendroglial subpopulations in Parkinson’s disease. Mol. Brain 16, 65 (2023).

37. Barba-Reyes, J. M. et al. Oligodendroglia vulnerability in the human dorsal striatum in Parkinson’s disease. Acta Neuropathol. (Berl.) 149, 46 (2025).

38. Adams, L., Song, M. K., Yuen, S., Tanaka, Y. & Kim, Y.-S. A single-nuclei paired multiomic analysis of the human midbrain reveals age- and Parkinson’s disease–associated glial changes. *Nat*. Aging 4, 364–378 (2024).

39. Osaki, Y. et al. Cross-sectional and longitudinal studies of three-dimensional stereotactic surface projection SPECT analysis in Parkinson’s disease. Mov. Disord. Off. J. Mov. Disord. Soc. 24, 1475–1480 (2009).

40. Goralski, T. M. et al. Spatial transcriptomics reveals molecular dysfunction associated with cortical Lewy pathology. Nat. Commun. 15, 2642 (2024).

41. Adler, C. H. et al. Unified Staging System for Lewy Body Disorders: Clinicopathologic Correlations and Comparison to Braak Staging. J. Neuropathol. Exp. Neurol. 78, 891–899 (2019).

42. Moore, J. et al. Longitudinal multi-omics in alpha-synuclein Drosophila model discriminates disease- from age-associated pathologies in Parkinson’s disease. Npj Park. Dis. 11, 46 (2025).

43. Brown, R. S. E. et al. Conditional Deletion of the Prolactin Receptor Reveals Functional Subpopulations of Dopamine Neurons in the Arcuate Nucleus of the Hypothalamus. J. Neurosci. 36, 9173–9185 (2016).

44. Kumasaka, N., Knights, A. J. & Gaffney, D. J. Fine-mapping cellular QTLs with RASQUAL and ATAC-seq. Nat. Genet. 48, 206–213 (2016).

45. Taylor-Weiner, A. et al. Scaling computational genomics to millions of individuals with GPUs. Genome Biol. 20, 228 (2019).

46. GTEx Consortium. The Genotype-Tissue Expression (GTEx) project. Nat. Genet. 45, 580–585 (2013).

47. Soutar, M. P. M. et al. Regulation of mitophagy by the NSL complex underlies genetic risk for Parkinson’s disease at 16q11.2 and MAPT H1 loci. Brain J. Neurol. 145, 4349–4367 (2022).

48. Hicks, A. R. et al. The non-specific lethal complex regulates genes and pathways genetically linked to Parkinson’s disease. Brain J. Neurol. 146, 4974–4987 (2023).

49. Smeland, O. B. et al. Genome-wide Association Analysis of Parkinson’s Disease and Schizophrenia Reveals Shared Genetic Architecture and Identifies Novel Risk Loci. Biol. Psychiatry 89, 227–235 (2021).

50. Pampari, A. et al. ChromBPNet: bias factorized, base-resolution deep learning models of chromatin accessibility reveal cis-regulatory sequence syntax, transcription factor footprints and regulatory variants. 2024.12.25.630221 Preprint at 10.1101/2024.12.25.630221 (2024).

51. Karczewski, K. J. et al. The mutational constraint spectrum quantified from variation in 141,456 humans. Nature 581, 434–443 (2020).

52. Nott, A. et al. Brain cell type-specific enhancer-promoter interactome maps and disease-risk association. Science 366, 1134–1139 (2019).

53. Mustache: multi-scale detection of chromatin loops from Hi-C and Micro-C maps using scale-space representation | Genome Biology | Full Text. https://genomebiology.biomedcentral.com/articles/10.1186/s13059-020-02167-0.

54. Activity-by-contact model of enhancer–promoter regulation from thousands of CRISPR perturbations | Nature Genetics. https://www.nature.com/articles/s41588-019-0538-0.

55. Granja, J. M. et al. ArchR is a scalable software package for integrative single-cell chromatin accessibility analysis. Nat. Genet. 53, 403–411 (2021).

56. Lee, S. The association of genetically controlled CpG methylation (cg158269415) of protein tyrosine phosphatase, receptor type N2 (PTPRN2) with childhood obesity. Sci. Rep. 9, 4855 (2019).

57. Yasui, T. et al. Insulin granule morphology and crinosome formation in mice lacking the pancreatic β cell-specific phogrin (PTPRN2) gene. Histochem. Cell Biol. 161, 223–238 (2024).

58. Ruiz-Martínez, J., Azcona, L. J., Bergareche, A., Martí-Massó, J. F. & Paisán-Ruiz, C. Whole-exome sequencing associates novel CSMD1 gene mutations with familial Parkinson disease. Neurol. Genet. 3, e177 (2017).

59. Lee, S., Wu, M. C. & Lin, X. Optimal tests for rare variant effects in sequencing association studies. Biostatistics 13, 762–775 (2012).

60. Zmuda, A. J. et al. A universal metabolite repair enzyme removes a strong inhibitor of the TCA cycle. Nat. Commun. 15, 846 (2024).

61. Madrer, N. et al. Pre-symptomatic Parkinson’s disease blood test quantifying repetitive sequence motifs in transfer RNA fragments. Nat. Aging 5, 868–882 (2025).

62. Hasan, N. & Gregg, R. G. Cone Synaptic Function is Modulated by the Leucine-Rich Repeat Adhesion Molecule LRFN2. eNeuro 11, (2024).

63. McMillan, K. J. et al. Sorting nexin-27 regulates AMPA receptor trafficking through the synaptic adhesion protein LRFN2. eLife 10, e59432 (2021).

64. Morimura, N. et al. Autism-like behaviours and enhanced memory formation and synaptic plasticity in Lrfn2/SALM1-deficient mice. Nat. Commun. 8, 15800 (2017).

65. Thevenon, J. et al. Heterozygous deletion of the LRFN2 gene is associated with working memory deficits. Eur. J. Hum. Genet. 24, 911–918 (2016).

66. Sekiyama, K. et al. Disease-Modifying Effect of Adiponectin in Model of α-Synucleinopathies. Ann. Clin. Transl. Neurol. 1, 479–489 (2014).

67. Wu, L.-J. et al. Genetic enhancement of trace fear memory and cingulate potentiation in mice overexpressing Ca2+/calmodulin-dependent protein kinase IV. Eur. J. Neurosci. 27, 1923–1932 (2008).

68. Alawneh, I., Amburgey, K., Gonorazky, H. & Gorodetsky, C. CAMK4-related Case of Hyperkinetic Movement Disorder. Mov. Disord. Clin. Pract. 10, 707–709 (2023).

69. Zech, M., et al. A unique de novo gain-of-function variant in CAMK4 associated with intellectual disability and hyperkinetic movement disorder. Cold Spring Harb. Mol. Case Stud. 4, a003293 (2018).

70. Albin, R. L., Young, A. B. & Penney, J. B. The functional anatomy of basal ganglia disorders. Trends Neurosci. 12, 366–375 (1989).

71. Surmeier, D. J., Ding, J., Day, M., Wang, Z. & Shen, W. D1 and D2 dopamine-receptor modulation of striatal glutamatergic signaling in striatal medium spiny neurons. Trends Neurosci. 30, 228–235 (2007).

72. Grandi, F. C., Modi, H., Kampman, L. & Corces, M. R. Chromatin accessibility profiling by ATAC-seq. Nat. Protoc. 17, 1518–1552 (2022).

73. Neavin, D. et al. Demuxafy: improvement in droplet assignment by integrating multiple single-cell demultiplexing and doublet detection methods. Genome Biol. 25, 94 (2024).

74. Bais, A. S. & Kostka, D. scds: computational annotation of doublets in single-cell RNA sequencing data. Bioinformatics 36, 1150–1158 (2020).

75. Enrichr: interactive and collaborative HTML5 gene list enrichment analysis tool - PubMed. https://pubmed.ncbi.nlm.nih.gov/23586463/.

76. propeller: testing for differences in cell type proportions in single cell data | Bioinformatics | Oxford Academic. https://academic.oup.com/bioinformatics/article/38/20/4720/6675456?login=false.

77. Gschwind, A. R. et al. An encyclopedia of enhancer-gene regulatory interactions in the human genome. 2023.11.09.563812 Preprint at 10.1101/2023.11.09.563812 (2023).

78. Pedersen, B. S. et al. Effective variant filtering and expected candidate variant yield in studies of rare human disease. NPJ Genomic Med. 6, 60 (2021).

